# EpiBinder: A Multimodal Framework for Cell-Type-Specific Prediction and Interpretation of Transcription Factor Binding

**DOI:** 10.64898/2026.07.06.736502

**Authors:** Ruben Solozabal, Albert Baichorov, Irina Miodownik, Tamir Avioz, Le Song, Marcos Matabuena, Martin Takáč, Ariel Afek

## Abstract

Transcription factor (TF) occupancy *in vivo* depends not only on the underlying DNA sequence but also on the local epigenetic environment, which varies across cell types and strongly influences whether sequence-encoded binding potential becomes functional. Here we present *EpiBinder*, a multimodal deep-learning framework for cell-type-specific prediction of TF binding that jointly models DNA sequence with base-resolution epigenetic information, including cytosine methylation from whole-genome bisulfite sequencing and chromatin accessibility from DNase I hypersensitivity data. Across multiple human cell lines, *EpiBinder* consistently outperforms strong sequence-only baselines, improving TF-binding prediction by up to 10% in area under the precision-recall curve. Beyond predictive performance, *EpiBinder* provides base-level attribution maps that enable systematic interrogation of regulatory context, including candidate methylation-sensitive loci, contextual motif dependencies, and putative TF-TF interactions. These results position *EpiBinder* as a practical framework for modeling and exploring the local regulatory grammar underlying cell-type-specific TF occupancy.

## 1. Introduction

### 1.1. Motivation and Problem Definition

Accurate prediction of transcription factor (TF) occupancy *in vivo* remains a central challenge in regulatory genomics. Although sequence-based deep learning models have substantially improved the prediction of TF-DNA interactions [2, 49, 23], motif information alone is often insufficient to explain binding in living cells, where local chromatin accessibility, DNA methylation, and cofactor context strongly shape whether a potential binding site becomes occupied [46, 27, 26]. As a result, many sequence-matched sites remain unbound, whereas bona fide binding can occur at weak or context-dependent sites, particularly in a cell-type-specific manner. These observations suggest that TF recognition *in vivo* is governed not only by sequence motifs, but by a broader local regulatory grammar that integrates sequence with epigenetic and contextual cues.

To address this gap, we developed *EpiBinder*, a multimodal deep-learning framework that integrates DNA sequence with base-resolution epigenetic features to model TF binding in a cell-type-aware manner. *EpiBinder* combines one-hot encoded sequence with cytosine methylation from whole-genome bisulfite sequencing in CpG, CHG, and CHH contexts, together with DNase-based chromatin accessibility, and is trained on TF-binding labels derived from ChIP-seq peak calls. By explicitly incorporating local epigenetic context, *EpiBinder* is designed not only to improve TF-binding prediction, but also to provide an interpretable framework for interrogating the context-dependent determinants of binding across cellular states.

In this work, we show that *EpiBinder* captures chromatin features that distinguish cellular contexts, an ability that is particularly important for modeling enhancer regulation, as enhancers are generally more cell-type-specific than promoters [42]. To learn these context-specific regulatory rules, we train independent models for the well-characterized human cell lines GM12878, K562, and HepG2. We further evaluate transfer to underrepresented cell types and apply the framework to downstream predictive and interpretability analyses to characterize the cell-type-specific regulatory dependencies learned by the model.

### 1.2. Related Work

Deep learning has emerged as a powerful class of computational techniques to enable genome-wide prediction of regulatory activity.

**Fig. 1:**
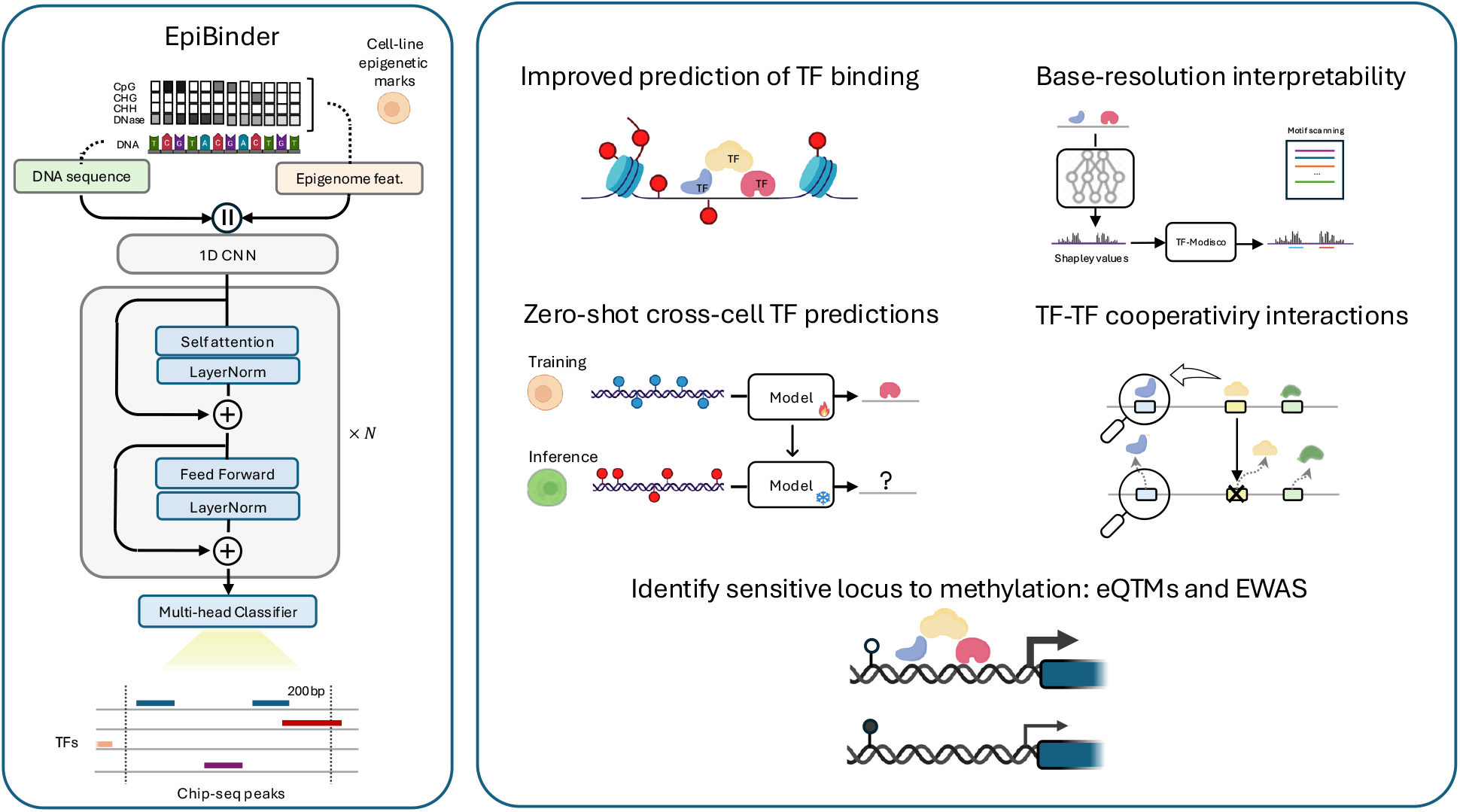
Overview of the EpiBinder framework and downstream tasks. **Left:** Multimodal architecture that encodes base-resolution DNA sequence alongside single base resolution epigenetic tracks (cytosine methylation and DNase I hypersensitivity) via parallel encoder streams, which are then fused to predict transcription factor binding. **Right:** Overview of the downstream tasks evaluated in this work, including zero-shot cross-cell TF prediction, detection of TF-TF cooperative interactions, and identification of methylation-sensitive loci via eQTMs and EWAS studies.

Early convolutional models such as DeepBind [2] showed that sequence-based neural networks can recover regulatory motifs and outperform classical machine learning and statistical methods in detecting TF binding. DeepSEA [49] extended this paradigm by scaling to large multitask prediction across diverse chromatin features and TFs. More recently, transformer-based architectures, including DNABERT [23, 51] and the Nucleotide Transformer [11], leverage pretraining models in large genomic corpora to learn transferable sequence representations. In this sense, models have evolved to capture a longer genomic context. Architectures such as Enformer [4], HyenaDNA [34], and AlphaGenome [5] expand receptive fields to hundreds of kilobases or more to model distal regulatory interactions. Although long-range context can help with tasks involving distal regulation, such as gene expression [4], TF binding in many cases is strongly shaped by local sequence features together with local chromatin and epigenetic state [34]. Motivated by this, rather than further extending the context length, we incorporate cell-type-specific epigenetic markers to better characterize the local binding environment. This distinction is especially relevant when the same genomic sequence is evaluated across different cellular contexts, where unchanged motif instances may nonetheless become functional or non-functional depending on cofactors, chromatin accessibility, and methylation state.

Parallel efforts have incorporated epigenetic information into the prediction of TF-binding. Methods such as SEMplMe [35] quantify methylation-dependent effects from ChIP-seq data, and linear or shallow models have incorporated CpG methylation as predictive covariates [21]. Deep learning models including DeepCpG [3], CpGPT [13] and MethylGPT [47] incorporate methylation signals, but are designed primarily to impute methylation or learn methylation-aware sequence representations, rather than to predict TF binding while explicitly integrating epigenetic information as regulatory context. These gaps suggest the need for multimodal predictors that combine the DNA sequence with local epigenetic information to achieve a cell–type-aware characterization.

### 1.3. Summary of Contributions

In this work, we introduce *EpiBinder* to address three related questions: (i) how much TF-binding prediction improves when local epigenetic context is modeled explicitly, (ii) whether such models can generalize across cellular contexts, and (iii) whether they can provide interpretable insight into the determinants of *in vivo* TF occupancy. Our main contributions are as follows:

1. **A multimodal framework for cell-type-specific TF-binding prediction**. We introduce *EpiBinder*, a base-resolution architecture that integrates DNA sequence, cytosine methylation, and chromatin accessibility to model TF binding in a cell-type-aware manner.
2. **Benchmark improvement over strong sequence-only baselines**. Across GM12878, HepG2, and K562, *EpiBinder* consistently improves TF-binding prediction relative to established sequence-only models and retains substantial predictive signal under cross-cell transfer.
3. **A framework for interrogating regulatory context**. Beyond prediction, *EpiBinder* enables base-level attribution analyses that prioritize candidate methylation-sensitive loci, contextual motif dependencies, and putative TF-TF interactions associated with cell-type-specific binding.

## 2. Materials and Methods

### 2.1. Data

The dataset used in this study corresponds to that described in [49], which comprises 919 chromatin features collected from ENCODE [14] and ROADMAP [8]. This corresponds to a well-characterized collection that has also been used to train TF modalities in Enformer [4] or AlphaGenome [5]. We particularized this dataset by selecting the 318 TF binding profiles from the three most represented cell lines: GM12878, K562 and HepG2, which together account for 45.9% of the ChIP-seq experiments in the dataset (Table S5). Binding occupancy is derived from processed ChIP-seq peak data binarized at 200 bp resolution, with the central 200 bp defining the target region and the 400 bp flanking sequences providing additional context, yielding a total 1,000 bp input window.

We augmented the dataset with epigenetic features corresponding to base-resolution methylation levels at the CpG, CHH and CHG sites derived from Whole-Genome bisulfite sequence (WGBS) and chromatin accessibility data from DNase I hypersensitivity assays (DNase-seq). Details of these datasets, including accession identifiers, are provided in the supplementary Table S6.

### 2.2. Model Description and Training Procedure

EpiBinder integrates DNA sequence and epigenetic information at single-base resolution within a multimodal deep learning framework to predict TF binding. Each genomic region is represented as a matrix of shape (*L*, 8), where *L* = 1000 bp denotes the length of the input sequence. The first four channels encode a one-hot representation of nucleotide bases (A, C, G, T), while the remaining four channels correspond to normalized epigenetic signals: cytosine methylation levels in CpG, CHG, and CHH contexts, plus a single continuous value representing DNase I hypersensitivity.

The architecture comprises a three-stage, one-dimensional convolutional backbone followed by a self-attention transformer encoder [44] and a feed-forward classification head. Each convolution stage applies a convolutional layer with ReLU activation, max-pooling, and dropout, progressively reducing the sequence dimensionality while projecting features into a shared latent space. The encoded representations after the transformer backbone are then flattened and passed through a linear classifier to produce TF-binding logits for each genomic position (detailed description in Table S7).

Separate models were trained for each cell line to explicitly capture cell-type–specific regulatory dynamics. Training was performed using binary cross-entropy loss optimized with the Adam optimizer, with early stopping. To prevent overfitting, layer normalization and dropout regularization were applied throughout the network.

### 2.3. Model Validation and Statistical Comparison

We train cell–line-specific *EpiBinder* models from scratch for the three most human characterized cell-lines in ENCODE: GM12878, K562, and HepG2. Training and testing sets were split by chromosomes to ensure non-overlap. In total, the model was trained with approximately 2.2 million training samples, with 227,512 held-out samples from chromosomes 8 and 9 reserved for testing. Under similar conditions, we compare our model against competitive sequence-only genomic predictors: DeepSEA [49], DNABERT [23], BigBird [48], and HyenaDNA [34]. We report auPRC rather than the more commonly used auROC, as auPRC offers a more informative measure for highly imbalanced datasets such as ChIP-seq, where positive peaks are relatively sparse (on average ∼17,000 peaks per TF and cell line).

In addition to improving predictive performance, our objective is to assess the capacity of the model to extrapolate across cellular contexts and to extract interpretable regulatory signals. Consequently, we complement our work with analyzes of cross-cell-line generalization and interpretability, including characterization of learned sequence patterns, sensitivity to epigenetic state, and context-dependent motif interactions. The evaluation is organized into four tasks, detailed in the following.

i. **Generalization ability to different cell-contexts**. Most available TF ChIP-seq datasets in ENCODE correspond to a small number of well-characterized cell lines, whereas for the majority of cells, only a limited set of TFs have been profiled (see ChIP-seq Atlas in [14]). Consequently, the ability to transfer knowledge from data-rich reference cell lines to under-characterized ones is a highly desirable property for predictive models. To assess this capability, we evaluated the zero-shot TF binding performance of *EpiBinder* in unseen cell lines, including A549 and H1-hESC. Specifically, we trained the model on a reference cell type and tested its out-of-domain (OOD) generalization on other cells with distinct epigenetic profiles. We quantified the performance drop between the in-domain (ID) and out-of-domain predictions using *Cohen’s d* (details in Supplementary S2). This analysis is informative not only when transfer succeeds, but also when it fails: TFs that are well predicted within each cell type yet exhibit poor cross-cell transfer provide evidence that the effective contextual determinants of occupancy can change across cellular environments even when the underlying genomic sequence is unchanged.
ii. **Interpreting EpiBinder through Pattern Discovery**. A central question motivating this study is whether genomic models genuinely recognize TF binding motifs or instead rely on indirect sequence-level signals to infer their presence. To investigate this, we systematically characterize the sequence patterns that *EpiBinder* learns when predicting TF binding events. Specifically, we interpret the model predictions using SHAP [32] and clustered the resulting importance scores using TF-MoDISco [43] to identify *seqlets*, or locally important subsequences, which were then consolidated into motif representations. This procedure is repeated for each TF, creating an *in silico library* of patterns that capture the learned binding preferences of the model. To evaluate the biological relevance of these discovered patterns, we performed a motif similarity analysis by computing pairwise *E-score* similarities between model-derived motifs and known TF motifs in the JASPAR database [40]. Furthermore, we use the resulting pairwise motif-TF association to quantify whether model-derived patterns map broadly to many reference motifs or instead concentrate on a single canonical motif. To this end, we introduce the Jaccard Overlap Score (JOS), which ranges from 0 when there are broad associations across diverse motifs, to 1 when there is a single dominant motif (formal definition in Supplementary S2.4).
iii. **Quantifying TF Cooperativity through targeted In Silico Perturbations**. While motif associations provide insight into sequence patterns linked to TF binding, they do not reveal whether these patterns arise from shared regulatory context, or functional dependencies between nearby factors. We therefore used targeted *in silico* perturbations to test whether local motifs systematically influence TF occupancy. Specifically, we constructed a dataset to analyze the effect of *in silico* randomization of co-binding motifs. We started with cell-type-specific enhancer annotations from EnhancerAtlas [17], where ChIP-seq peaks were used to identify candidate binding regions. We refined these broad ChIP-seq peaks to motif-level sites using FIMO [19] and enumerated motif pairs with centers within ±500bp. For each pair, we quantified interaction effects by querying the model before and after masking the *putative interacting TF* motif through nucleotide randomization, while keeping all other inputs fixed. The resulting change in predicted binding probability relative to baseline was then used to quantify the dependency between the motif pair: positive values indicate competition, such that disrupting the putative interacting motif increases predicted binding, whereas negative values indicate cooperativity, such that disrupting the motif decreases predicted binding. Secondly, we particularize the study on the effects of TF interactions on enhancer *SORT1* in HepG2 cell, for which MPRA data are available [25]. To evaluate the prediction capabilities of the model, we performed an *in-silico* saturation mutagenesis. At each position within the enhancer, we introduced all possible single-nucleotide substitutions and quantified the average change in the predicted probability of TF-binding across all factors. This approach provides a base-resolution map of the general model’s sensitivity at each binding location and enables us to study context-dependent TF interactions within the enhancer.
iv. **Assess the Impact of DNA Methylation on TF Binding**. Finally, we evaluate the regulatory impact of cytosine methylation on TF binding. To this end, we performed *in silico* genome-wide methylation experiments in silico using the trained *EpiBinder* model to identify CpG sites within a ±500bp genomic window where methylation significantly influences predicted TF binding. For each site, the change in predicted TF-binding probability between the original and modified inputs was used to quantify the local methylation sensitivity. The high-sensitivity CpGs were then cross-referenced with the Illumina EPIC 850k array to identify the sites commonly profiled in EWAS studies. Finally, functional enrichment analysis was conducted to determine whether these loci overlapped promoters, enhancers, or other regulatory regions, using a two-sided Fisher’s exact test with Benjamini–Hochberg correction for multiple comparisons.

## 3. Results

We next assess the performance of *EpiBinder* and explore the regulatory signals captured by the model. We first quantify the predictive gains for TF-binding prediction across cell lines (Section 3.1). We then assess generalization performance when transferring across cell-lines (Section 3.2). We then characterize the binding determinants captured by the model (Section 3.3). We probe context-dependent TF–TF interactions through targeted *in silico* perturbations to quantify cooperativity and competitive effects (Section 3.4). Finally, we use the model’s epigenetic sensitivity to identify CpG methylation loci associated with regulatory activity (Section 3.5).

### 3.1. *EpiBinder* improves TF-binding prediction across all evaluated cell lines relative to strong sequence-only baselines

The results of the statistical associations are described in Table 1. Overall, *EpiBinder* improves TF-binding prediction compared to current genomic models [34], DNABERT [23], BigBird [48], and DeepSEA [49]. While Table 1 reports mean auPRC values averaged across all evaluated TFs within each cell line, Figure 2 complements this by showing the distribution of auPRC values across all TFs evaluated for each cell-line, thereby highlighting the variability in performance between TFs. This heterogeneity is partly explained by data availability, Figure 2b shows a positive association between the number of available ChIP-seq peaks and the model’s performance, indicating that TFs with fewer binding events remain more challenging to predict.

**Table 1.** Mean auPRC per cell-line grouped for Transcription factor binding (TF) and Polymerase (Pol). The same testing conditions on chromosomes 8 and 9 are preserved from [49].

| MODEL | LEN | GM12878 |  | HepG2 |  | K562 |  |
| --- | --- | --- | --- | --- | --- | --- | --- |
|  |  | TF | Pol | TF | Pol | TF | Pol |
| DNA sequence |  |  |  |  |  |  |  |
| DeepSea | 1kbp | 26.1 | 35.7 | 29.3 | 36.1 | 25.8 | 31.8 |
| HyenaDNA | 1kbp | 26.9 | 37.6 | 30.5 | 33.3 | 26.5 | 32.3 |
| DNABERT | 512bp | 26.7 | 35.3 | 29.1 | 35.7 | 26.0 | 31.3 |
| BigBird | 8kbp | 27.4 | 34.8 | 31.7 | 34.4 | 26.7 | 30.9 |
| DNA sequence + epigenetics |  |  |  |  |  |  |  |
| EpiBinder | 1kbp | **39.1** | **48.4** | **43.1** | **45.4** | **41.1** | **44.7** |

**Fig. 2:**
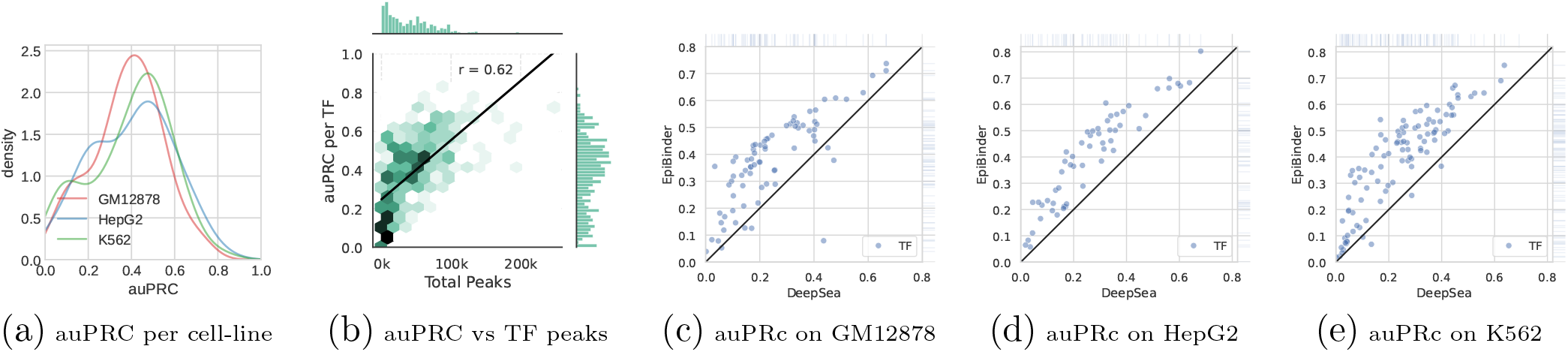
Results on TF-binding prediction. **(a)** Density distributions of per-cell auPRC for EpiBinder. **(b)** Scatter plot of binding prediction performance versus training data availability. The x-axis shows the number of ChIP-seq peaks available for each TF, and the y-axis shows the corresponding in-domain auPRC. In black, the best-fit linear regression. **(c–e)** Scatterplots of per-TF auPRC comparing DeepSEA (x-axis) vs EpiBinder (y-axis) predictions on GM12878, HepG2, and K562 cell-lines.

Although previous sequence-based models exhibit broadly similar performance despite architectural differences, EpiBinder delivers consistent and substantial improvements across all cell lines evaluated (see Table 1). Relative to the foundational DeepSEA model, EpiBinder achieves an average auPRC increase of **+14.0%** for transcription factor (TF) binding and **+11.6%** for RNA polymerase prediction. These gains are particularly notable given that each *EpiBinder* model is trained on considerably fewer examples, as each model is restricted to TF ChIP-seq data from a single cell line, underscoring the advantage of explicitly modeling cell-type–specific epigenetic information. As shown in Figures 2c–e, improvements are observed for nearly all TFs, with the largest gains occurring for those with intermediate baseline performance (auPRC ≈ 0.2–0.6). Even for TFs with low baseline performance (auPRC *<* 0.2), *EpiBinder* yields consistent improvements despite sparse ChIP-seq coverage.

Beyond TF binding, *EpiBinder* also enhances histone-mark prediction, achieving an average auPRC gain of **+18.2%** compared with DeepSEA (Table 2).

**Table 2.** Histone mark predictions on K562 cell-line reported as auPRC. Testing conditions are preserved from [49].

|  | DeepSea<br>—40M— | HyenaDNA<br>—7M— | DNABERT<br>—86M— | BigBird<br>—110M— | EpiBinder<br>—46M— |
| --- | --- | --- | --- | --- | --- |
| H2AZ | 0.45 | 0.43 | 0.42 | 0.44 | **0.65** |
| H3K27ac | 0.46 | 0.44 | 0.42 | 0.46 | **0.70** |
| H3K27me3 | 0.09 | 0.08 | 0.08 | 0.09 | **0.25** |
| H3K36me3 | 0.14 | 0.13 | 0.15 | 0.15 | **0.42** |
| H3K4me1 | 0.33 | 0.34 | 0.31 | 0.35 | **0.67** |
| H3K4me2 | 0.56 | 0.54 | 0.52 | 0.58 | **0.78** |
| H3K4me3 | 0.58 | 0.56 | 0.54 | 0.60 | **0.78** |
| H3K79me2 | 0.33 | 0.32 | 0.29 | 0.35 | **0.47** |
| H3K9ac | 0.52 | 0.50 | 0.49 | 0.53 | **0.75** |
| H3K9me1 | 0.04 | 0.04 | 0.04 | 0.04 | **0.08** |
| H3K9me3 | 0.06 | 0.06 | 0.10 | 0.07 | **0.12** |
| H4K20me1 | 0.13 | 0.13 | 0.12 | 0.13 | **0.21** |

### 3.2. Zero-Shot Prediction of TF Binding Across Cell Lines

Figure 3 summarizes zero-shot TF-binding performance across cell lines. For each pair of cell lines, and for all TFs with available data in both, we computed the auPRC obtained from zero-shot predictions. As observed, incorporating epigenetic markers markedly enhances cross-cell generalization. We also quantified the performance drop between in-domain (ID) and out-of-domain predictions using *Cohen’s d* (GM12878: *d* ≈ 0.8–1.1; HepG2: *d* ≈ 0.5–0.8; K562: *d* ≈ 0.7–1.1; see Table 4). These performance gaps indicate measurable degradation under cross-cell transfer, but also show that *EpiBinder* retains substantial predictive signal across distinct cellular contexts. Additionally, Figure 3b illustrates the correlation between OOD/ID predictions for a representative subset of 29 TFs profiled in GM12878, HepG2, and K562 (full results in Supplementary Table S9). For all cell-lines, OOD auPRC values show a strong correlation with their ID counterparts (average Pearson’s *r* = 0.83), confirming that the model captures a substantial fraction of transcription factor binding variability through shared epigenomic features.

**Fig. 3:**
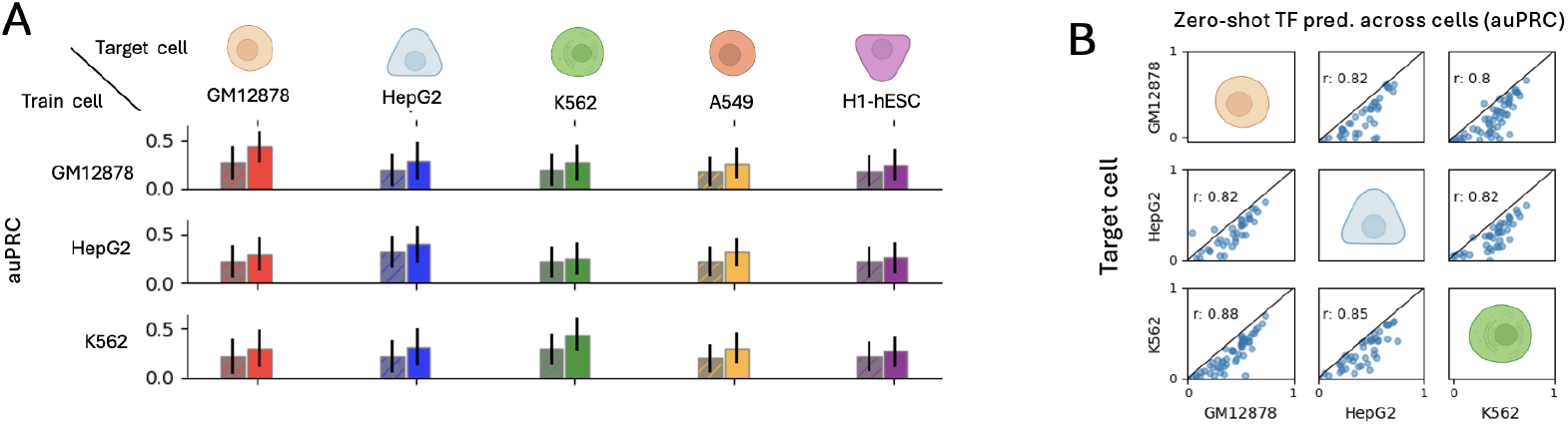
Zero-shot binding predictions across cell-lines. **(a)** Zero-shot TF binding prediction across different cell-lines for the model, including the epigenetic features (in color) or shadowed for the model relying only on the nucleotide sequence. **(b)** Correlation on the zero-shot predictions for a subset of TF concurrently available in the studied cell-lines.

Interestingly, for TFs with few binding events, zero-shot transfer from a related, data-rich cell line can occasionally outperform the corresponding ID model. Table 3 summarizes representative cases where this occurs. For example, ZNF274 in HepG2 exhibits only 1,090 ChIP-seq peaks, yielding an in-domain auPRC of 0.00; whereas a model trained on GM12878, which provides 9,358 peaks (8.5× more data), achieves an out-of-domain auPRC of 0.30, corresponding to a +30% absolute improvement. As illustrated in Figure 2e, the number of training peaks is positively correlated with predictive performance. Thus, transferring information from well-characterized cell lines effectively compensates for limited in-domain profiles, enabling improved TF-binding predictions even under different cell conditions.

**Table 3.** Top TFs ranked by absolute improvement in auPRC when transferring from in-domain (ID) to out-of-domain (OOD) predictions. For each TF and target cell line, we report the number of ChIP-seq peaks in the ID and OOD datasets, the OOD/ID peak ratio, the corresponding auPRC values, and the absolute auPRC gains.

| TF | CELL-LINE | ID PEAKS | OOD PEAKS | RATIO OOD/ID PEAKS | ID auPRC | OOD auPRC | auPRC GAIN |
| --- | --- | --- | --- | --- | --- | --- | --- |
| ZNF274 | HepG2 | 1090 | 9358 | $\times 8.5$ | 0.00 | 0.30 | +30% |
| CHD2 | HepG2 | 14700 | 54416 | $\times 3.7$ | 0.23 | 0.40 | +17% |
| Mxi1 | K562 | 21104 | 71904 | $\times 3.4$ | 0.38 | 0.52 | +14% |
| RFX5 | K562 | 7174 | 18654 | $\times 2.6$ | 0.10 | 0.21 | +11% |
| TR4 | K562 | 2280 | 8968 | $\times 3.9$ | 0.04 | 0.15 | +11% |

**Table 4.** Generalization performance of models trained on three different cell lines (GM12878, HepG2, and K562) when evaluated across both their in–domain (id) and out–of–domain (ood) cell conditions. For each training cell (“in–domain”), we report the mean auPRC performance (*µ*) and standard deviation (*σ*) on that cell as well as on each external test cell line. The column Δ shows the absolute drop in mean performance relative to the “out-of–domain” result (*µ*_id_ *− µ*_ood_), *R* gives the relative percentage drop (Δ*/µ*_id_ *×* 100%), and Cohen’s *d* quantifies the standardized effect size of that gap.

| Cell Line | $\mu$ | $\sigma$ | $\Delta = \mu_{id} - \mu$ | $R = \frac{\Delta}{\mu_{id}}$ (%) | Cohen's $d$ |
| --- | --- | --- | --- | --- | --- |
| **GM12878<sup>(ID)</sup>** | 0.43 | 0.16 | — | — | — |
| HepG2 | 0.29 | 0.19 | 0.15 | 33.8 | 0.82 |
| K562 | 0.27 | 0.18 | 0.17 | 38.1 | 0.91 |
| A549 | 0.26 | 0.16 | 0.17 | 39.4 | 1.04 |
| H1 | 0.24 | 0.17 | 0.19 | 44.0 | 1.10 |
| **HepG2<sup>(ID)</sup>** | 0.40 | 0.18 | — | — | — |
| GM12878 | 0.30 | 0.17 | 0.10 | 25.3 | 0.57 |
| K562 | 0.26 | 0.17 | 0.14 | 35.6 | 0.81 |
| A549 | 0.32 | 0.15 | 0.08 | 20.9 | 0.50 |
| H1 | 0.26 | 0.16 | 0.14 | 35.4 | 0.82 |
| **K562<sup>(ID)</sup>** | 0.44 | 0.18 | — | — | — |
| GM12878 | 0.30 | 0.20 | 0.14 | 32.5 | 0.76 |
| HepG2 | 0.32 | 0.19 | 0.13 | 28.6 | 0.68 |
| A549 | 0.30 | 0.15 | 0.14 | 31.2 | 0.83 |
| H1 | 0.26 | 0.15 | 0.18 | 40.2 | 1.07 |

### 3.3. EpiBinder Exhibits Broad Predictive Associations for Identifying TF Binding

To investigate the sequence and contextual determinants of binding learned by *EpiBinder*, we compared model-derived motifs with experimentally validated TF motifs from the JASPAR database.

The resulting map (Fig. 4a) shows that *EpiBinder* captures a broad spectrum of motif–TF associations across most factors. At the same time, the model reveals substantial diversity in pattern coverage across TFs (Fig. 4b). Whereas a small subset of TFs is dominated by a single canonical motif, most TFs exhibit a heterogeneous mixture of secondary patterns. We quantify this tendency using the JOS score (Fig. 4c), which distinguishes narrowly focused motif mappings from broader, multi-motif association profiles.

**Fig. 4:**
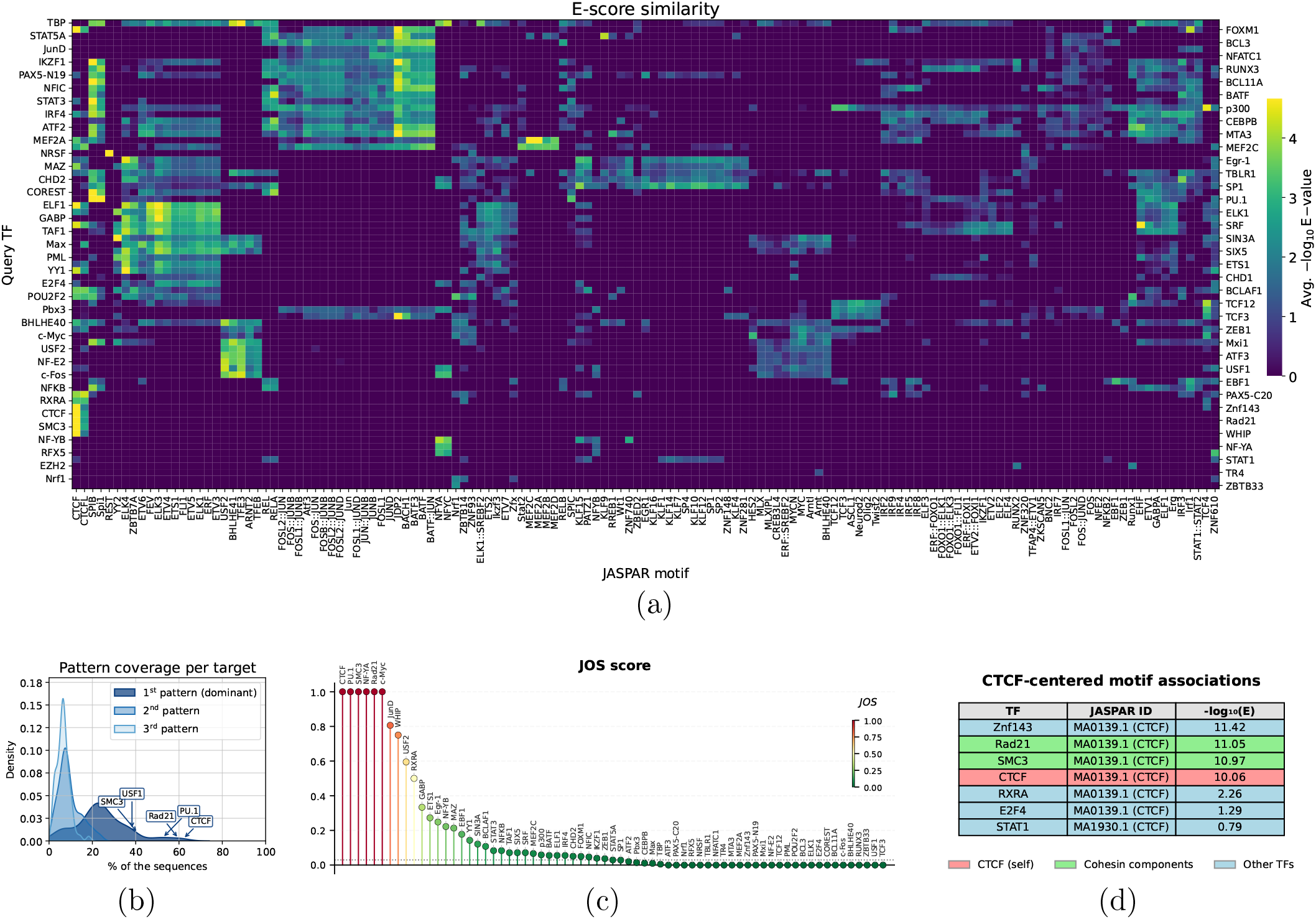
Model-derived patterns reveal widespread contextual associations (GM12878 cell-line). **(a)** Motif similarity between model-derived patterns and the JASPAR database reported as the Tomtom similarity score (*−* log_10_ of the E-value). Bright colors correspond to stronger similarity (lower E-values), while darker colors indicate low similarity. **(b)** Density estimates of per-target for the top *k* = 3 patterns discovered across the targets. The dominant pattern shows the highest coverage. Annotations show the TFs with the highest presence of the dominant pattern. **(c)** Jaccard Overlap Score (JOS) summarizing diversity in the association for each TF. The score ranges from 0 (patterns exhibit wide associations with diverse motifs) to 1 (patterns converge on a single canonical motif). **(d)** Network representation of factors having in common CTCF as the dominant pattern. The diagram highlights the cohesin components (RAD21, SMC3) that present CTCF motifs as the dominant pattern.

More broadly, the motif association map reveals distinct modes of binding organization. Some factors are associated almost exclusively with their own canonical motif, consistent with localization driven primarily by direct sequence recognition. Others are strongly influenced by motifs associated with additional regulators, suggesting that their occupancy is shaped by recruitment, co-localization, or broader regulatory context. A third group displays broad multi-motif association profiles, indicating that their predicted binding depends on a richer combination of local sequence and contextual signals rather than on a single dominant determinant.

To understand what these different association profiles may reflect biologically, we next examined systems in which the roles of direct sequence recognition and context-dependent recruitment are already well characterized. The CTCF–cohesin system provides a particularly informative example, as it includes factors with distinct modes of genomic localization that have been extensively characterized experimentally [29, 12, 9]. CTCF itself shows one of the highest JOS values, and the dominant motif identified by *EpiBinder* closely matches its canonical JASPAR motif, consistent with localization driven largely by direct sequence recognition. By contrast, the cohesin components RAD21 and SMC3 also exhibit high JOS values, but their dominant associated motif is CTCF rather than a factor-specific motif. This is consistent with the fact that cohesin does not recognize DNA through a dedicated sequence motif. Instead, cohesin is recruited to and stabilized at specific genomic sites through interactions with sequence-anchored factors such as CTCF at chromatin boundaries and loop domains [12, 9]. Thus, the recovery of a CTCF-centered signature for RAD21 and SMC3 suggests that *EpiBinder* captures this indirect recruitment mechanism.

This behavior is not restricted to cohesin. Across several factors, the model shows dominant associations with motifs of other strong regulators rather than with their own canonical motif (Supplementary Table S10), suggesting that *EpiBinder* captures broader dependency structure within the regulatory landscape. In particular, the CTCF motif contributes strongly to the prediction of multiple additional factors (Fig. 4d), consistent with the role of CTCF as an architectural anchor in chromatin organization. Related CTCF-centered associations have been reported for ZNF143 [6, 50] and RXRA [10, 45]. Extending this analysis across cell types (Supplementary Table S11), we find that the composition of CTCF-centered associations varies markedly across cellular contexts. In HepG2, the strongest associations involve FOXA-family motifs, consistent with the role of FOXA proteins as hepatic pioneer factors and with evidence that CTCF can cooperate with lineage-determining pioneer TFs to organize regulatory hubs [22, 31]. In K562, the corresponding associations instead highlight hematopoietic regulators, including PU.1 and C/EBP-family factors, in agreement with their established roles in specifying lineage-defining enhancer programs [20, 38, 15]. Overall, these results suggest that *EpiBinder* captures a mixture of direct architectural dependencies, lineage-specific cooperative occupancy, and indirect chromatin-context correlations.

At the opposite extreme, factors such as TBP and p300 exhibit broad multi-motif association profiles, indicating that their predicted occupancy is shaped by a wider combination of local regulatory features rather than by a single dominant sequence determinant. Despite this plethora of patterns, in some cases the behavior remains relatively transferable. As TBP with mean auPRC decreasing only from 0.44 in domain to 0.33 out of domain (see Table S9), suggesting that the features learned for TBP are shared across the 3 evaluated cellular contexts. In contrast, p300 shows much weaker transferability, with mean auPRC dropping from 0.39 in domain to 0.13 out of domain (supplementary Table S9), indicating strong cell specificity.

However, motif associations alone do not distinguish whether these patterns reflect direct recognition, correlated context, or directional dependencies between nearby regulators. We therefore next used targeted in silico perturbations to test whether local motifs exert systematic effects on predicted TF occupancy.

### 3.4. Quantifying Context-Dependent TF Cooperativity Through In Silico Motif Perturbation

To probe candidate context-dependent TF-TF interactions, we performed targeted *in silico* perturbations of nearby motifs and measured the resulting effect on the TF of interest. For each binding locus detected genome-wide across the enhancer regions, we infer candidate cooperative or competitive dependencies between motifs through systematic perturbation. Specifically, we first computed the predicted binding probability of the TF of interest under the unperturbed sequence. We then systematically randomize the motif of putative interacting partners (*perturbed TF*), and recalculate the resulting binding probabilities for the TF of interest. This procedure enables quantification of cooperative and competitive interactions between the TF of interest and its putative interacting partners.

The results are presented in Figure 5a, where each entry represents the average change in the predicted binding relative to the baseline. Strong negative values indicate that removal of the partner motif substantially reduces the predicted binding of the motif of interest. In contrast, positive values denote competitive interaction, where the loss of a partner motif enhances the binding. In the figure, we collect the most influential TFs, which consistently correspond to architectural or pioneer-like regulators. In GM12878, CTCF and AP-1 components (FOSL2::JUND, JUND) dominated; the loss of CTCF broadly weakened adjacent motifs, reflecting its established role in chromatin insulation and loop anchoring. In K562, the most impactful TFs were GATA1, GATA2, MAX, and MAX::MYC, which correspond to lineage-defining erythroid regulators with pioneering activity that promote co-binding. In HepG2, consistent with [37], members of the FOX family (FOXA2, FOXA3, FOXK1, FOXK2, and FOXO1) exhibited the strongest effects, reflecting their established roles as pioneer factors (FOXA) and as components of enhancer-associated complexes that shape cooperative occupancy [39, 22].

**Fig. 5:**
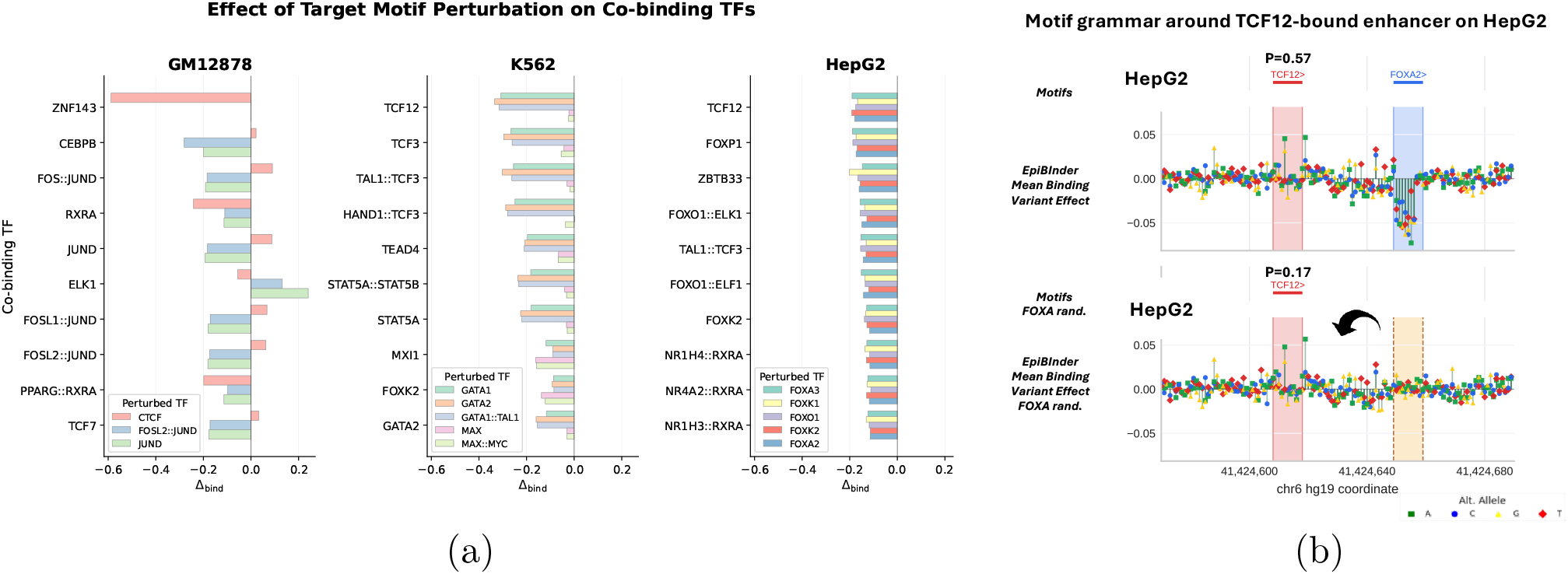
In silico perturbation experiments reveal cell-type–specific TF-TF cooperativity interactions. **(a)** Barplots correspond to the effect of randomizing a partner motif and analyzing its effect on the co-binding TF of interest. We report the signed absolute value of the randomization effect relative to the baseline prediction across different cell-lines. (**b**) Example of TCF12-bound enhancer on HepG2 at chr6:41424570–41424690 with a nearby FOX-associated motif. *EpiBinder* -derived position-wise mutagenesis profile detects that when predicting TCF12 (red) occupancy, the model assigns strong sensitivity to the neighboring FOX-associated motif (blue). Randomizing the FOX motif significantly reduces the predicted TCF12 binding probability.

Among the targets, TCF12 emerges as one of the most prominent examples of context-dependent factors. The perturbation analysis reveals that its predicted occupancy depends on different local partners in each cellular context. In HepG2, perturbation of nearby FOX-associated motifs substantially decreases predicted TCF12 binding (avg. Δ_bind_ = −0.31), whereas in K562, TCF12 is most sensitive to nearby GATA-associated motifs (avg. Δ_bind_ = −0.19). To illustrate this behavior, we selected representative enhancer examples with cell-type-specific TCF12 predictions. In Fig. 5b, *EpiBinder* predicts TCF12 binding at a HepG2-specific enhancer, but not in K562. At this locus, the local mutagenesis profile highlights a nearby FOX-associated motif as a major contributor to the HepG2 prediction, and randomization of this motif reduces the predicted TCF12 binding probability from 0.57 to 0.17 (Δ_bind_ = −0.40). Conversely, at the K562-specific locus highlighted in Supplementary Fig. 10, the model predicts TCF12 binding in K562 but not in HepG2, with sensitivity concentrated around nearby GATA-associated motifs. Together, these examples illustrate how *EpiBinder* relies on the cell-type-specific regulatory partners in the local chromatin context to predict binding.

On the other hand, the ELK1–JUND/AP-1 pair was identified as a competitive interaction (Fig. 5a). Across GM12878 enhancer loci, randomization of nearby JUND/AP-1 motifs recurrently increased predicted ELK1 binding, with an average effect of Δ_bind_ = +0.22 (a representative locus is shown in Supplementary Fig. 9). This positive perturbation effect suggests that, in GM12878, JUND/AP-1 motifs can create an antagonistic local context for ELK1 occupancy rather than a cooperative one, which is consistent with known context dependences in the literature [36, 41, 33].

We then applied the same position-wise perturbation framework to the SORT1 enhancer in HepG2, where experimental MPRA measurements are actually available. The resulting *in silico* mutagenesis profile is shown in Figure 6a. At each position in the enhancer, we introduced all possible single-nucleotide substitutions and quantified the resulting change in the model’s average TF-binding profile. Although *EpiBinder* does not directly model gene expression, these position-wise predicted effects showed a significant correspondence with the experimentally measured impact on gene expression from MPRA, with a correlation of *r* = 0.393. Notably, the model accurately recovered 4 of the 7 transcription factors previously reported [1] to bind and regulate the enhancer, including key hepatocyte-specific regulators such as FOXA1, HNF4G, USF1, and SP5. The remaining factors, including SOX6, NFYA, and KLF5, exhibited weaker binding signals in the model.

**Fig. 6:**
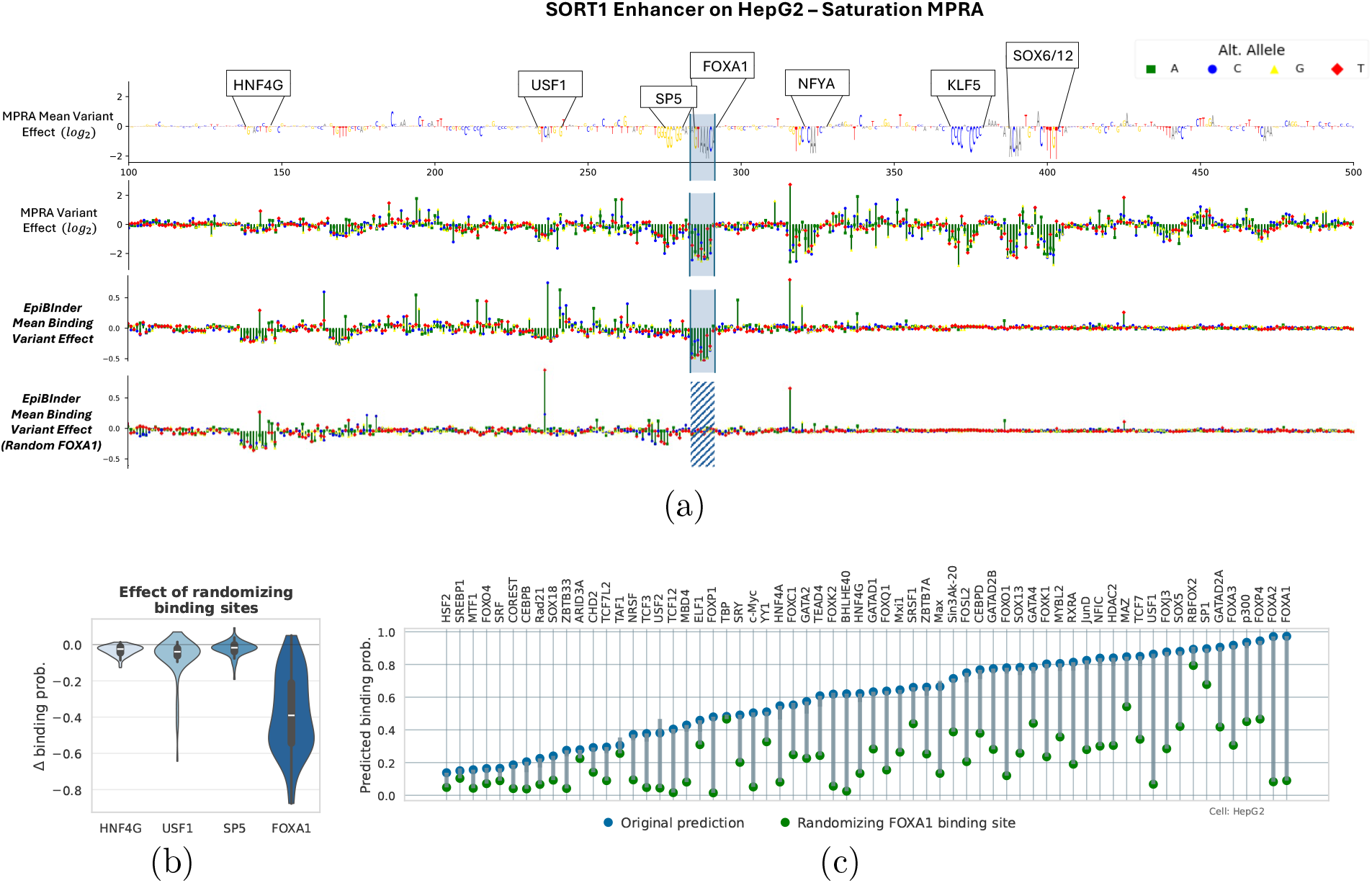
Regulatory activity at the *SORT1* enhancer in HepG2. (**a**) Correlation between predicted nucleotide-level regulatory effects from *in silico* saturation mutagenesis and experimentally measured MPRA activity for the *SORT1* enhancer. Each point represents a single-nucleotide substitution, and the observed correlation (*r ≈* 0.39). The top TFs predicted by the model (FOXA1, HNF4G, USF1 and SP5) coincide with known regulators of hepatic lipid metabolism. (**b**) Effect of motif randomization on the transcription factor co-binding probabilities predicted by the model. (**c**) Particularized predictions before (blue) and after (green) randomization of the FOXA1 binding site. Loss of FOXA1 binding results in a pronounced reduction in co-binding probabilities for neighboring transcription factors, highlighting its function as a pioneer factor.

We also assessed cooperativity among the binding sites identified by *EpiBinder* within the enhancer. For each of the four detected binding sites, we study the effect of motif randomization in the cofactor predictions. Particularly, we quantify the average change in prediction heads in Fig. 6b. Perturbation of HNF4G, USF1, and SP5 produced only moderate effects in the binding cofactors, suggesting that these motifs make comparatively limited independent contributions to the predicted binding landscape. In contrast, randomization of the FOXA1 motif led to a pronounced bulk-effect reduction across most binding prediction heads, indicating that the predictive contribution of neighboring motifs is substantially strengthened when FOXA1 is present in the enhancer. Figure 6c provides a detailed view of the FOXA1 perturbation results, illustrating a significant decrease in predicted binding probabilities not only for FOXA1 itself but also for multiple co-located TFs. Together, these results suggest that motif presence alone is insufficient to accurately predict TF occupancy, and that reliable prediction requires a well-characterized local cellular context, which in the case of SORT1 enhancer in HepG2 cells, appears to be strongly shaped by FOXA1.

### 3.5. In Silico Identification of CpG_m_ Sites Associated with Regulatory Activity

We next applied *EpiBinder* to identify genome-wide CpG_m_ loci where DNA methylation plays a critical role in transcriptional regulation. For each 1,000bp window across the whole dataset (genome-wide), we compared the predicted TF-binding score before and after setting its methylation level to zero, flagging as “high-sensitivity” those sites with Δ_*bind*_ values within the top quantile. Across GM12878, K562, and HepG2, we recovered two major classes of loci: (i) *demethylation-enhanced* sites, where loss of methylation substantially increases predicted TF affinity, and (ii) *demethylation-repressed* sites, which are rarer and predominantly observed in K562 and HepG2.

To refine these predictions, we integrated the identified CpG_m_ sites with data from the Illumina EPIC BeadChip, which profiles DNA methylation at over 850,000 CpG sites across the human genome (Fig. 7a). Cross-referencing these data enabled us to investigate the potential regulatory relevance of such positions. In particular, we first focused on sites where removal of the methyl mark led to a pronounced *in silico* increase in predicted TF occupancy. Although these regions show no TF to bind in the native cell line, we speculate that such CpG_m_ loci may function as methylation-controlled regulatory regions, potentially enabling the binding of regulatory TFs. If that is the case, we speculate they may serve as active regulatory elements in other cellular contexts.

**Fig. 7:**
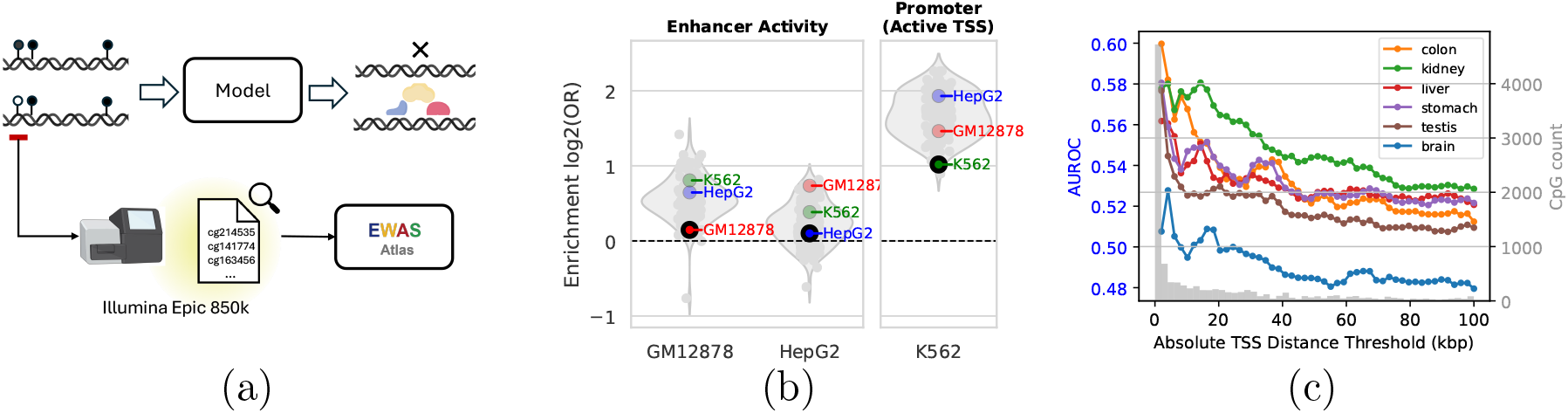
Identification of CpG methylation–sensitive loci. (**a**) CpG positions showing strong methylation sensitivity were cross-referenced with the Illumina EPIC 850k array and annotated according to their regulatory context. (**b**) Methylation-sensitive CpG_m_ sites discovered within inactive regions of GM12878 and HepG2 are nonetheless markedly enriched for enhancer-associated chromatin states across numerous Roadmap cell types, whereas CpG_m_ sites identified in K562 exhibit preferential enrichment within promoter-associated states. (**c**) Association between predicted methylation effects on TF binding and expression-associated CpGs (mQTL) from the EWAS database. The auROC quantifies the discriminative power of the model-predicted change in CpG methylation levels to gene expression across tissues and genomic distances.

To further support this hypothesis, we assessed the enrichment of the detected CpG_m_-sensitive sites within different chromatin states using data from the Roadmap Epigenomics project. For each chromatin state, enrichment was calculated as the ratio between the proportion of CpG probes overlapping that feature and the corresponding background frequency across all Roadmap samples. This analysis revealed that CpG sites predicted by *EpiBinder* to enhance TF binding in GM12878 and HepG2 are effectively associated with enhancer regions across multiple Readmap cell types, whereas those detected in K562 show stronger enrichment in promoter-associated chromatin states (Fig. 7b). These findings suggest that methylation-sensitive loci detected by the model contribute to distinct regulatory activity depending on cellular context.

Finally, to evaluate whether the CpG_m_ loci identified by *EpiBinder* correspond to expression-associated CpGs, we examined their overlap with methylation quantitative trait loci (mQTLs) curated in the EWAS Atlas [28]. This resource provides correlation statistics linking the methylation levels of CpG sites to the expression of nearby genes. We specifically tested whether CpG_m_ sites predicted by the model to recruit or repel transcription factors have the discriminative power to distinguish positively and negatively correlated CpG–gene pairs. We use the area under the receiver operating characteristic (auROC) metric across multiple tissues and genomic distances to evaluate these capabilities. The resulting profiles (Fig. 7c) show that the model predictions are most informative for promoter-proximal CpGs, with performance gradually declining at greater genomic distances. Notably, tissues corresponding to the cell-line contexts used for *EpiBinder* training (e.g., hematopoietic and hepatic) exhibit the strongest signals, suggesting that predicted TF modulation at CpG_m_ loci captures biologically meaningful mechanisms linking DNA methylation to transcriptional regulation.

## 4. Discussion

In this work, we present *EpiBinder*, a base-resolution multimodal framework that integrates DNA sequence with single-base-resolution epigenetic signals to model TF binding in a cell– type-aware manner. *EpiBinder* consistently outperforms strong sequence-only baselines, yielding higher auPRC while operating on a compact 1 kbp input window. These results question the prevailing emphasis on extending receptive fields to hundreds of kilobases or to the megabase scale [4, 5]. Although long-context models are motivated by distal regulatory interactions, our experiments suggest that increasing sequence span alone offers limited benefit for TF-binding prediction beyond a few kilobases [34]. Instead, improved characterization of *local* regulatory context emerges as the dominant driver of performance improvements.

Beyond improving accuracy, *EpiBinder* retains substantial predictive power under cellular-context shifts, enabling to transfer the knowledge from cell to cell. This is especially useful in settings where TF-binding data is sparse or when analyzing cell-types that remain poorly characterized. Mechanistically, *EpiBinder* provides a lens for interrogating what deep genomic models learn about regulation: we show that it captures context-dependent binding logic, including motif-pair dependencies that are consistent with both cooperative and competitive effects. Finally, we connect epigenetic variation to downstream function by linking methylation-sensitive predictions to expression-associated CpG–gene relationships and quantitative methylation trait loci (mQTLs).

Several limitations motivate future work. First, our evaluation is restricted to a small number of well-characterized cell lines, largely due to limited availability of highly profiled cell-types with extensive TF ChIP-seq collected. Deep genomic models often benefit from joint learning in which many TFs are predicted by the same model, which enables shared regulatory representation and improves sample efficiency. In addition, base-resolution interpretability remains computationally demanding: the shapley-value attribution utilized [32] is expensive on a genomic scale, and therefore our motif discovery and interpretability analyzes were derived from a limited subset of sites (1,000 binding events). More scalable attribution methods and broader sampling will be needed to assess the generality of these mechanistic conclusions.

As multi-omic resources continue to expand, multimodal approaches such as *EpiBinder* should become increasingly effective for predictive and mechanistic modeling of non-coding regulatory regions, enabling stronger cross-cellular transfer and more informative hypotheses about determinants of TF occupancy. Future extensions that incorporate additional modalities, including histone modifications and 3D chromatin topology [16, 24], could further advance this framework toward unified models that jointly capture sequence, chromatin state, and genome organization.

## 5. Supplementary data

Supplementary Data are available at NAR Genomics and Bioinformatics Online.

## 7. Data availability

The data underlying this article were derived from publicly available datasets from ENCODE and the NIH Roadmap Epigenomics Project, as described in the Materials and Methods. Accession identifiers for the WGBS and DNase-seq datasets used in this study are provided in Supplementary Table S6. The version 0.0.1 of the EpiBinder software used for this article is permanently archived in Zenodo at https://zenodo.org/records/21966803.

## Supplementary Material

### S1. Model design and training data

We describe details on the data set used in this work. Table 5 reports, for each of the six most characterized ENCODE cell-lines, the number of TFs with ChIP-seq experiments and their fraction of the total TF in the collection, motivating our emphasis on cell types with dense TF coverage. Table 6 then lists the ENCODE accession identifiers for the epigenetic modalities used in this study.

**Table 5.**
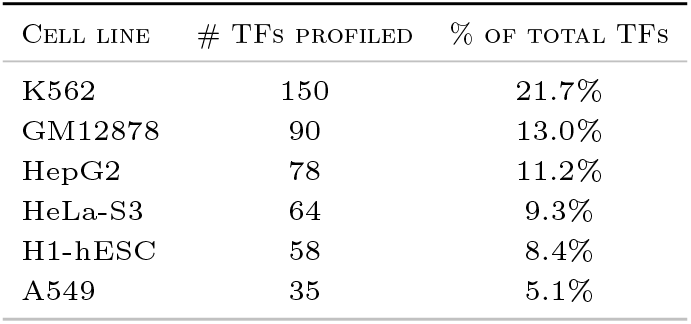
Number and percentage of TFs profiled in the ENCODE dataset for the six most extensively characterized human cell lines. These cell lines collectively account for the majority of available ChIP-seq experiments used in [49].

**Table 6.** ENCODE accession identifiers for epigenetic datasets used in this study: Whole-Genome Bisulfite Sequencing (WGBS) and DNase I hypersensitivity assays used to model chromatin accessibility.

| CELL |  | WGBS | DNASE-SEQ |
| --- | --- | --- | --- |
| GM12878 | CpG | ENCFF570TIL | ENCFF264NMW |
|  | CHH | ENCFF187KAK |  |
|  | CHG | ENCFF910HOG |  |
| HepG2 | CpG | ENCFF817LMT | ENCFF867UYB |
|  | CHH | ENCFF158FZM |  |
|  | CHG | ENCFF101UQI |  |
| K562 | CpG | ENCFF660IHA | ENCFF352SET |
|  | CHH | ENCFF294NMQ |  |
|  | CHG | ENCFF571AGF |  |
| A549 | CpG | ENCFF948WVD | ENCFF674WCO |
|  | CHH | ENCFF589SGV |  |
|  | CHG | ENCFF461YYK |  |
| H1-hESC | CpG | ENCFF434CNG | ENCFF233CHA |
|  | CHH | ENCFF036NWK |  |
|  | CHG | ENCFF780ECA |  |

The details about the model are as follows. EpiBinder architecture maps a length-*L* = 1000 genomic window to multi-label TF-binding predictions by jointly modeling DNA sequence and epigenetic context. For each mini-batch of size *B*, the input is encoded as the channel-wise concatenation

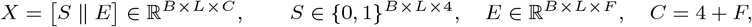

where *S* denotes the one-hot nucleotide encoding and *E* stacks the normalized epigenetic tracks (channel definitions in Table 7). A three-stage 1D convolutional encoder extracts hierarchical representations, 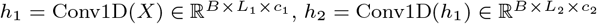, and 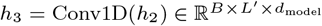. The resulting sequence of latent vectors is processed by a transformer encoder, *Z* = Transformer 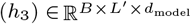, whose layers use multi-head self-attention with scaled dot-product attention Attn(*Q, K, V*) = softmax 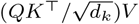.

**Table 7.** Summary of the layers configuration and output dimensions for *EpiBinder* architecture.

| Stage | Layer(s) & Configuration | Output Shape |
| --- | --- | --- |
| Input | One-hot nucleotide + 4 normalized epigenetic | $(B, L, 8)$ |
| Conv Layer 1 | Conv1d(8→320, k=8, padding=4) | $(B, \lfloor L/4 \rfloor, 320)$ |
| Conv Layer 2 | Conv1d(320→480, k=8, padding=4) | $(B, \lfloor L/16 \rfloor, 480)$ |
| Conv Layer 3 | Conv1d(480→$d_{\text{model}}$, k=8, padding=4) | $(B, L', d_{\text{model}})$ |
| Positional Embedding | Embedding(num_embeddings=$L'$, embedding_dim=$d_{\text{model}}$) | $(B, L', d_{\text{model}})$ |
| Transformer Encoder | $N \times \text{EncoderLayer}(d_{\text{model}}, n_{\text{heads}})$ | $(B, L', d_{\text{model}})$ |
| Flatten | — | $(B, L' \times d_{\text{model}})$ |
| Classifier MLP | $\text{MLP}(L' \times d_{\text{model}} \rightarrow d_{\text{clf}} \rightarrow d_{\text{clf}} \rightarrow n_{\text{labels}})$ | $(B, n_{\text{labels}})$ |

Finally, *Z* is flattened, 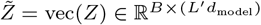, and passed to an MLP classifier 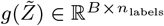; an element-wise sigmoid yields TF-binding probabilities for each label.

### S2. Methods

#### S2.1. Pattern Collection and Motif Similarity Analysis

To characterize the sequence and epigenetic determinants driving the model’s TF-binding predictions, we computed the base-resolution attributions and summarized them into motifs. For each cell line, we randomly sampled *N* = 1000 genomic regulatory regions we verified the TF of interest effectively binds. For each region, we formulate the input construction *X* = [*S*|*E*], and we use a trained model to output a prediction score *ŷ* = *g*_*θ*_(*X*), which we used to derive the feature attributions.

Particularly, we compute base-resolution importance scores using DeepSHAP [32], obtaining attribution values for all input channels (nucleotide and epigenetic) at each genomic position. Let *s*_*i,c*_ denote the attribution for channel *c* ∈ 1, …, *C* at position *i* ∈ 1, …, *L*. We summarized these channel-wise attributions into a single position-level importance score by summing the absolute attribution mass from (i) the nucleotide channels and (ii) the epigenetic channels,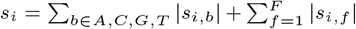 . To extract interpretable sequence patterns from the resulting profiles, we applied TF-MoDISco [43]. TF-MoDISco identifies short high-signal subsequences (“seqlets”) by clustering windows with consistently large attribution mass (based on |*s*_*i*_|), and then aligns and clusters seqlets using similarity of contribution-weighted representations to produce a non-redundant set of motif-like patterns.

Finally, to assess biol3ogical relevance, we compared the resulting collection of patterns to the JASPAR database of curated motifs [40] using Tomtom [7]. Tomtom computes similarity statistics between model-derived patterns and reference motifs (measured as e-score), enabling us to link discovered patterns to canonical TF binding sites or, in some cases, broader regulatory contexts.

#### S2.2. Construction of the TF–TF Cooperativity Dataset

To study context-dependent TF interactions, we constructed a TF–TF cooperativity dataset that integrates local DNA sequence with cell-type–specific regulatory context. We began with enhancer annotations from EnhancerAtlas [17] and restricted analyses to enhancers assigned to each of the three studied cell lines. Within these enhancers, we used ChIP-seq peak calls from [49] to define candidate TF-bound regions. Because ChIP-seq peaks are typically broad and do not localize motif instances precisely, we refined peak coordinates by scanning for motif matches with FIMO. For the set of all enhancers, we searched for motif instances consistent with the ChIP-seq–labeled TF; when FIMO reported motif complexes, we retained a match if at least one component corresponded to the annotated TF.

Using the resulting set of localized motif instances, we enumerated all TF pairs (*t*_*s*_, *t*_*n*_) occurring within a genomic distance of at most 500 bp, where *t*_*s*_ denotes the *source* TF and *t*_*n*_ its *target* neighbor. For each pair, we extracted a length-*L* = 1000 bp window centered on the source motif, encoded it as *X* = [*S*|*E*], and queried the model to obtain a baseline binding probability for the source TF, *p*_0_ = *g*_*θ*_(*X*). To quantify interdependence between the two motifs, we performed *in silico* randomization experiments in which the nucleotide sequence spanning the neighboring motif was replaced by randomized bases (using *r* = 3 independent random seeds), yielding perturbed inputs *X*^(*r*)^ and corresponding predictions *p*_*r*_ = *g*_*θ*_(*X*^(*r*)^), while all other sequence and epigenetic features in the window were held fixed. We defined the dependency score as Δ*p* = E*r*[*p*_*r*_ − *p*_0_], measuring how perturbing the neighbor motif alters predicted binding at the source motif under an identical regulatory context.

#### S2.3. Standardized difference between OOD/ID performance

To quantify how strongly model performance degrades under cross-cell generalization, we computed Cohen’s *d* between in-domain (ID) and out-of-domain (OOD) evaluations using AUPRC. For each ordered pair of cell lines (*c*_id_, *c*_ood_), we restricted analysis to the set of TF labels shared by both cell lines, *T* (*c*_id_, *c*_ood_). For each *t* ∈ *T* (*c*_id_, *c*_ood_), we computed the per-TF AUPRC under ID evaluation held-out data from *c*_id_, and the per-TF AUPRC under OOD evaluation by evaluating the same model on data from *c*_ood_. We then summarized these distributions across shared TFs by their means and standard deviations, (*µ*_id_, *σ*_id_) and (*µ*_ood_, *σ*_ood_), and computed

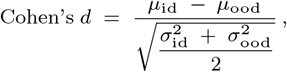

where the denominator is the pooled standard deviation, yielding a scale-free measure of the AUPRC gap between ID and OOD conditions. Larger *d* indicates a greater discrepancy between ID and OOD performance. In this study, Cohen’s *d* provides an interpretable summary of cross-cell generalization, enabling direct comparisons of how well *EpiBinder* retains predictive power under distributional shifts in cell-specific regulatory context.

#### S2.4. Jaccard Overlap Score (JOS)

In this paper, we introduce the Jaccard Overlap Score (JOS) to quantify whether the model-derived patterns for a given TF reflect a wide associations or instead collapse onto a small number of canonical motifs. For each transcription factor *t*, TF-MoDISco clusters high-attribution “seqlets” (high-importance subsequences extracted from the model interpretation) into a set of motif-like patterns, which we denote by *P*_*t*_ = {*p*_1_, …, *p*_|*·*|_}. Each pattern *p* ∈ *P*_*t*_ can be viewed as a cluster prototype summarizing a family of similar seqlets and is represented by an aggregated motif. To relate these learned patterns to known motifs, we run Tomtom against JASPAR and define J (*p*) as the set of significant JASPAR motif matches returned for pattern *p* (using *q*-value ≤ 0.05). We then quantify agreement between all patterns using the Jaccard index, and define JOS as one minus the mean pairwise overlap:

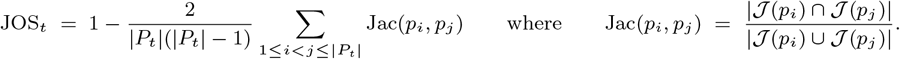

Here Jac(*p*_*i*_, *p*_*j*_) is high when the two pattern-clusters map to the same (or highly overlapping) subset of JASPAR motifs and low when they map to different subsets. Therefore, JOS_*t*_ ∈ [0, 1]: values near 0 indicate strong agreement among pattern-clusters (many patterns share the same JASPAR matches, consistent with collapse onto a restricted motif set), whereas values near 1 indicate weak agreement (indicating diverse motif associations).

### S3. Tools

We used well-established tools for motif discovery and motif matching. In particular, the tools and configuration parameters used are as follows:

**TF-MoDISco** [43]: applied to cluster high-importance subsequences (“seqlets”) identified from attribution scores into motif-like patterns. We use the default false discovery rate (FDR ≤ 0.05).

**MEME Suite / Tomtom** [7]: employed to measure similarity between model-derived motifs and known binding motifs from the JASPAR database [40]. Tomtom computes statistical similarity scores and provides motif annotations. We use it, setting a minimum overlap of 5 bp, and a significance threshold *q* ≤ 0.01.

**FIMO** [19]: used to refine broad ChIP-seq peaks by scanning for precise motif instances, ensuring alignment between experimentally observed binding and sequence-level motif coordinates. We applied a significance threshold of *p* ≤ 1 × 10^*−*4^.

### S4. Reproducing efforts on the Chromatin profile prediction task

To ensure a fair comparison in our study, we reproduced representative baselines on the chromatin profile prediction task using publicly available implementations and retraining recipes described in the original works. Crucially, all models were trained and evaluated using the *same* dataset construction and the *same* train/validation/test splits, so that differences in performance reflect model and training choices. We report in Table 8 mean auROC across three assay categories—TF binding, DNase I hypersensitivity (DHS), and histone modifications (HM)—to match the standard evaluation setting on the original works.

**Table 8.**
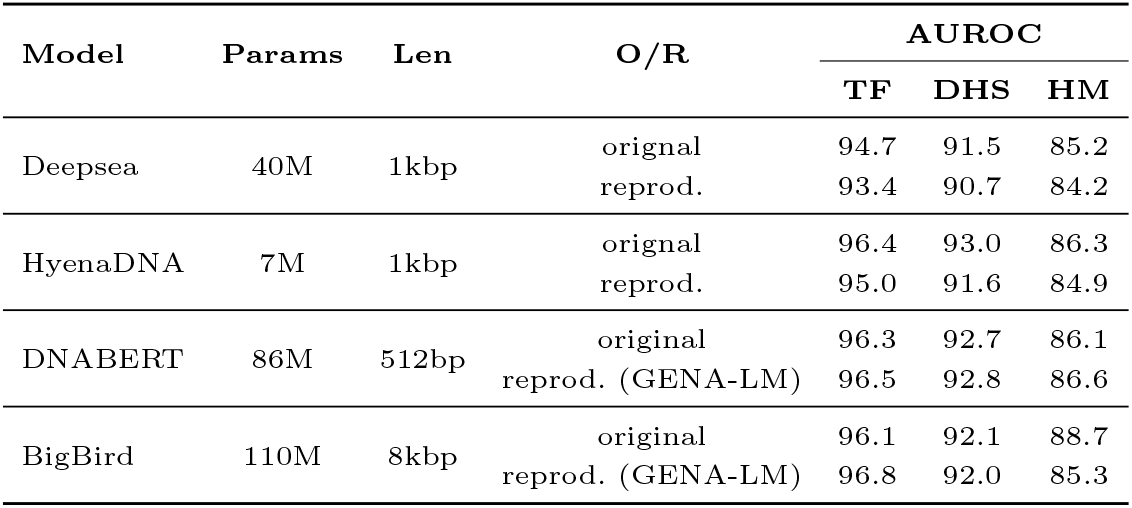
Performance comparison between reported in the original work and reproduced efforts on the Chromatin profile prediction task. Mean area under the ROC curve (auROC), averaged across three assay types: transcription factor binding profiles (TF), DNase I–hypersensitive sites (DHS), and histone modifications (HM).

| Model | Params | Len | O/R | AUROC |  |  |
| --- | --- | --- | --- | --- | --- | --- |
|  |  |  |  | TF | DHS | HM |
| Deepsea | 40M | 1kbp | original | 94.7 | 91.5 | 85.2 |
|  |  |  | reprod. | 93.4 | 90.7 | 84.2 |
| HyenaDNA | 7M | 1kbp | original | 96.4 | 93.0 | 86.3 |
|  |  |  | reprod. | 95.0 | 91.6 | 84.9 |
| DNABERT | 86M | 512bp | original | 96.3 | 92.7 | 86.1 |
|  |  |  | reprod. (GENA-LM) | 96.5 | 92.8 | 86.6 |
| BigBird | 110M | 8kbp | original | 96.1 | 92.1 | 88.7 |
|  |  |  | reprod. (GENA-LM) | 96.8 | 92.0 | 85.3 |

Details on the reproduded models are as follows:

- **DeepSEA**. We reproduce the dataset construction and retrain the model following [30].
- **HyenaDNA**. We fine-tune a HyenaDNA-7M model to reproduce the results obtained in the original work [34]. For the downstream task of chromatin profile prediction, we utilize a pre-trained Hyena encoder in combination with sequence-level pooling and a fully connected decoder to perform multilabel sequence classification. In the original work, two variants with sequence lengths of 1k and 8k were fine-tuned. As reported in the paper, the 1k model outperforms the 8k model in predicting short-range tasks such as transcription factor (TF) binding. Therefore, we reproduce this setting for performance comparison.
- **GENA-LM** We finetune a Gena-LMs Bert and BigBrid model using the downstream code provided in [18].

### S5. Additional results

We present complementary results that expand on analyses discussed in the main text. Table 9 summarizes zero-shot transfer performance across cell lines for the subset of TFs shared between GM12878, HepG2 and K562 cell lines (Section 3.2). Figure 8 reports motif enrichment at demethylation-sensitive loci identified in the in silico methylation perturbation analysis (Section 3.5). Table 10 provides additional examples of motif co-associations beyond the CTCF case study highlighted in Section 3.3).

**Table 9.** Comparison of zero-shot auPRC performance across different cell-lines. The top label indicates the cell-line the model was trained on, whereas the bottom label indicates the cell-line being tested. *in-domain* refers to the scenario in which the model is both trained and tested on the same cell-line, included here for comparison. As observed, some cases show an improvement when transferring prediction from one cell-line to another.

| Source cell | GM12878 |  |  | HepG2 |  |  | K562 |  |  |
| --- | --- | --- | --- | --- | --- | --- | --- | --- | --- |
| Target cell | *in-domain* | HepG2 | K562 | *in-domain* | GM12878 | K562 | *in-domain* | HepG2 | GM12878 |
| SMC3 | **0.73** | 0.60 | 0.69 | **0.67** | 0.65 | 0.60 | **0.73** | 0.59 | 0.70 |
| CTCF | **0.65** | 0.64 | 0.55 | **0.72** | 0.54 | 0.53 | **0.65** | 0.63 | 0.55 |
| Rad21 | **0.63** | 0.61 | 0.55 | **0.65** | 0.57 | 0.56 | **0.64** | 0.59 | 0.54 |
| GABP | 0.61 | **0.64** | 0.47 | **0.65** | 0.57 | 0.46 | 0.58 | **0.59** | 0.50 |
| TAF1 | 0.51 | **0.56** | 0.49 | **0.70** | 0.39 | 0.47 | 0.58 | **0.62** | 0.45 |
| MAZ | **0.57** | 0.44 | 0.52 | **0.52** | 0.50 | 0.49 | **0.67** | 0.26 | 0.39 |
| ELF1 | **0.62** | 0.36 | 0.41 | **0.50** | 0.42 | 0.41 | **0.53** | 0.44 | 0.49 |
| NRSF | **0.25** | **0.25** | 0.15 | **0.48** | 0.34 | 0.25 | **0.23** | 0.09 | 0.03 |
| Nrf1 | **0.51** | 0.32 | 0.37 | **0.42** | 0.28 | 0.36 | **0.41** | 0.40 | 0.35 |
| USF1 | **0.50** | 0.36 | 0.35 | **0.56** | 0.37 | 0.38 | **0.48** | 0.41 | 0.35 |
| Max | **0.50** | 0.37 | 0.39 | **0.50** | 0.45 | 0.41 | **0.61** | 0.42 | 0.38 |
| YY1 | **0.34** | 0.20 | 0.17 | **0.52** | 0.17 | 0.31 | 0.46 | **0.50** | 0.19 |
| USF2 | **0.53** | 0.39 | **0.53** | **0.57** | 0.40 | 0.49 | **0.57** | 0.42 | 0.41 |
| Mxi1 | **0.51** | 0.48 | 0.26 | **0.58** | 0.45 | 0.26 | 0.38 | **0.52** | 0.47 |
| CEBPB | **0.30** | 0.03 | 0.04 | **0.53** | 0.04 | 0.22 | **0.48** | 0.31 | 0.07 |
| TBP | **0.42** | 0.37 | 0.27 | **0.44** | 0.35 | 0.32 | **0.48** | 0.41 | 0.31 |
| JunD | **0.34** | 0.05 | 0.08 | **0.54** | 0.04 | 0.23 | **0.57** | 0.23 | 0.05 |
| SP1 | **0.54** | 0.18 | 0.34 | **0.45** | 0.28 | 0.15 | **0.45** | 0.23 | 0.43 |
| c-Myc | **0.37** | 0.10 | 0.21 | 0.26 | 0.20 | **0.31** | **0.50** | 0.25 | 0.15 |
| ATF3 | **0.30** | 0.28 | 0.19 | **0.35** | 0.23 | 0.28 | **0.39** | 0.24 | 0.12 |
| CHD2 | **0.51** | 0.15 | 0.26 | 0.23 | **0.40** | 0.27 | 0.34 | 0.18 | **0.38** |
| BHLHE40 | **0.43** | 0.21 | 0.25 | **0.31** | 0.29 | 0.24 | **0.42** | 0.24 | 0.30 |
| p300 | **0.45** | 0.09 | 0.08 | **0.38** | 0.21 | 0.10 | **0.36** | 0.13 | 0.19 |
| RFX5 | **0.28** | 0.20 | 0.05 | **0.29** | 0.19 | 0.07 | 0.10 | **0.21** | 0.13 |
| ZNF274 | 0.03 | 0.00 | **0.10** | 0.00 | **0.30** | 0.00 | **0.39** | 0.00 | 0.00 |
| COREST | **0.20** | 0.13 | 0.10 | **0.22** | 0.11 | 0.13 | **0.39** | 0.09 | 0.06 |
| TR4 | **0.11** | 0.06 | 0.01 | **0.20** | 0.02 | 0.04 | 0.04 | **0.15** | 0.02 |
| SRF | **0.20** | 0.04 | 0.09 | 0.06 | **0.10** | 0.08 | **0.14** | 0.05 | **0.14** |
| ZBTB33 | **0.13** | 0.08 | 0.05 | **0.20** | 0.10 | 0.05 | **0.13** | 0.06 | 0.07 |

**Table 10.**
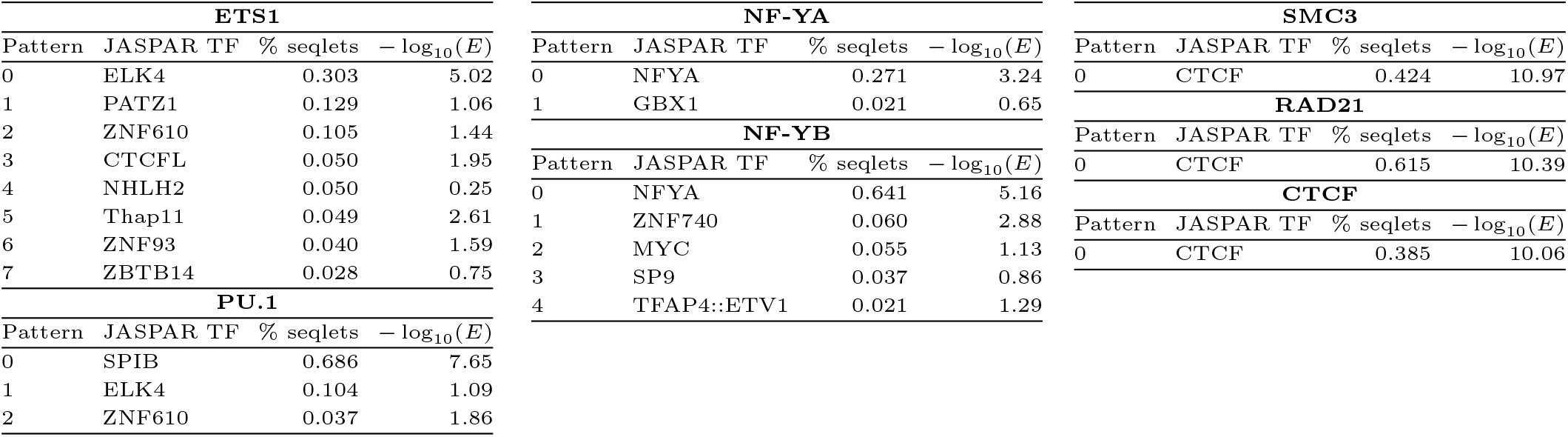
Motif co-associations. Discovered sequence patterns were matched to JASPAR motifs; we report the top match, % seqlets assigned to each pattern, and *−* log_10_(*E*). Dominant matches recover expected identities and reveal co-associations where the model links a TF’s predictive signal to correlated motifs from related factors (e.g., ETS1, PU.1*→*ETS-family) or to architectural anchors (SMC3/RAD21*→*CTCF). Such co-associations may reflect a motif confounding problem, as they are interpreted as model-level dependencies rather than true interactions.

**Table 11.**
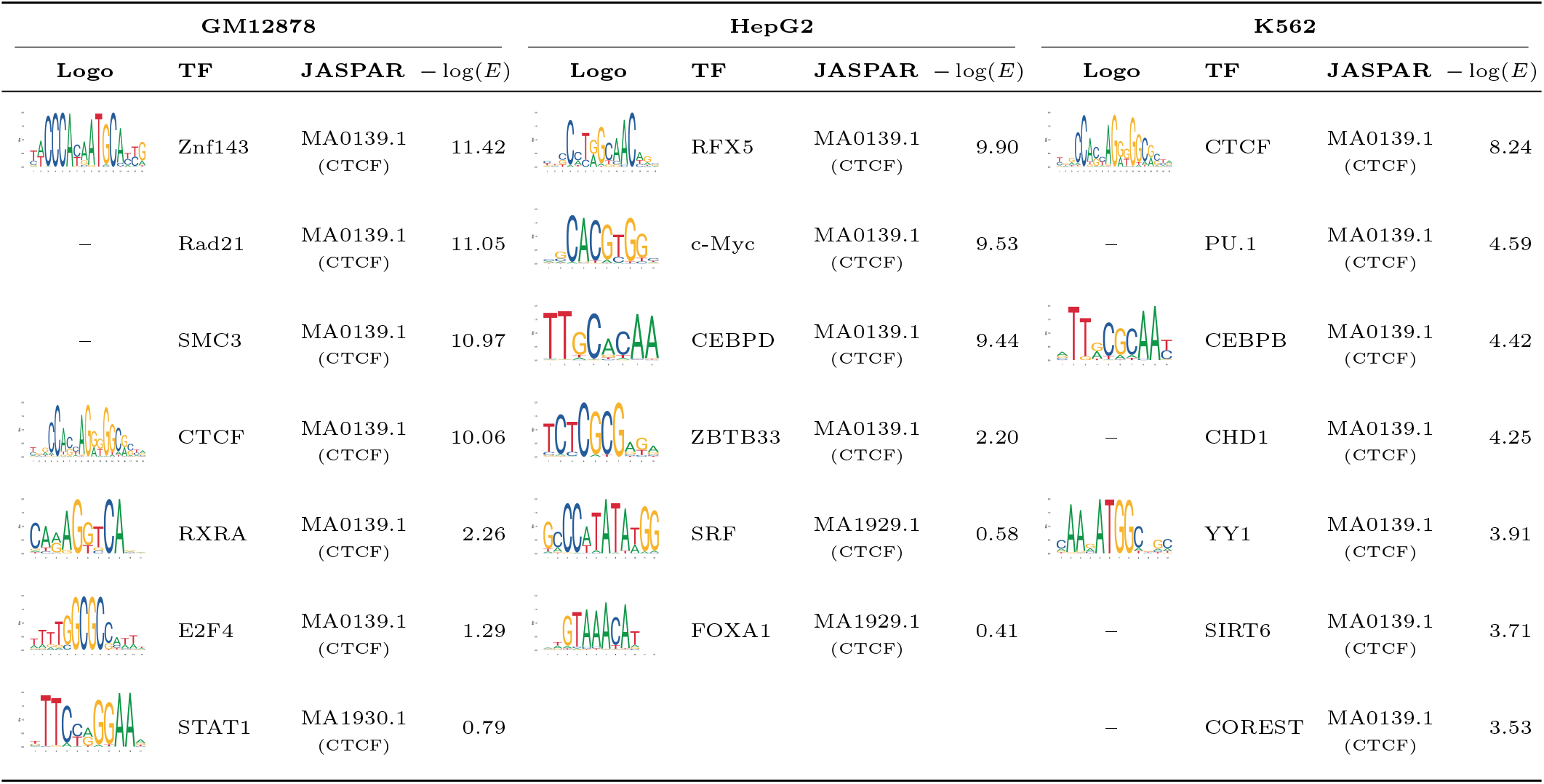
Top model-inferred transcription factor associations with CTCF-centered motifs across the GM12878, HepG2, and K562 cell lines. For each cell context, the table reports the associated factor, the corresponding motif logo, the matched JASPAR motif annotation, and the enrichment score (*−* log(*E*)). The differences across cell lines highlight the cell-specific regulatory context of CTCF binding detected by *EpiBinder*. Motif logos are shown when a corresponding JASPAR-derived logo file was available; otherwise, a hyphen is displayed.

| GM12878 |  |  |  | HepG2 |  |  |  | K562 |  |  |  |
| --- | --- | --- | --- | --- | --- | --- | --- | --- | --- | --- | --- |
| Logo | TF | JASPAR | $-\log(E)$ | Logo | TF | JASPAR | $-\log(E)$ | Logo | TF | JASPAR | $-\log(E)$ |
|  | Znf143 | MA0139.1<br>(CTCF) | 11.42 |  | RFX5 | MA0139.1<br>(CTCF) | 9.90 |  | CTCF | MA0139.1<br>(CTCF) | 8.24 |
| — | Rad21 | MA0139.1<br>(CTCF) | 11.05 |  | c-Myc | MA0139.1<br>(CTCF) | 9.53 | — | PU.1 | MA0139.1<br>(CTCF) | 4.59 |
| — | SMC3 | MA0139.1<br>(CTCF) | 10.97 |  | CEBPD | MA0139.1<br>(CTCF) | 9.44 |  | CEBPB | MA0139.1<br>(CTCF) | 4.42 |
|  | CTCF | MA0139.1<br>(CTCF) | 10.06 |  | ZBTB33 | MA0139.1<br>(CTCF) | 2.20 | — | CHD1 | MA0139.1<br>(CTCF) | 4.25 |
|  | RXRA | MA0139.1<br>(CTCF) | 2.26 |  | SRF | MA1929.1<br>(CTCF) | 0.58 |  | YY1 | MA0139.1<br>(CTCF) | 3.91 |
|  | E2F4 | MA0139.1<br>(CTCF) | 1.29 |  | FOXA1 | MA1929.1<br>(CTCF) | 0.41 | — | SIRT6 | MA0139.1<br>(CTCF) | 3.71 |
|  | STAT1 | MA1930.1<br>(CTCF) | 0.79 | — | — | — | — | — | COREST | MA0139.1<br>(CTCF) | 3.53 |

**Table 12.** Top model-inferred transcription factor associations with FOXA1-centered motifs in HepG2 and GATA2-centered motifs in K562. The prominent association of TCF12 with these motif-centered contexts is consistent with the perturbation analysis in Section 3.4, in which randomization of nearby FOXA and GATA motifs strongly reduced predicted TCF12 binding. Motif logos are shown when a corresponding JASPAR-derived logo file was available; otherwise, a hyphen is displayed.

| FOXA1 (HepG2) |  |  |  | GATA2 (K562) |  |  |  |
| --- | --- | --- | --- | --- | --- | --- | --- |
| Logo | TF | JASPAR | $-\log_{10}(E)$ | Logo | TF | JASPAR | $-\log_{10}(E)$ |
|  | Nrf1 | MA0148.4<br>(FOXA1) | 2.20 | — | TBLR1 | MA0036.3<br>(GATA2) | 2.80 |
|  | ZBTB7A | MA0148.4<br>(FOXA1) | 1.40 | — | SETDB1 | MA0036.3<br>(GATA2) | 2.70 |
|  | MYBL2 | MA0148.4<br>(FOXA1) | 1.40 | — | HMGN3 | MA0036.3<br>(GATA2) | 2.60 |
| — | MBD4 | MA0148.4<br>(FOXA1) | 1.30 | — | HDAC6 | MA0036.3<br>(GATA2) | 2.20 |
|  | **TCF12** | MA0148.4<br>(FOXA1) | **1.30** | — | TAL1 | MA0036.3<br>(GATA2) | 2.10 |
|  | JunD | MA0148.4<br>(FOXA1) | 1.20 | — | BRF2 | MA0036.3<br>(GATA2) | 2.00 |
|  | FOXA1 | MA0148.4<br>(FOXA1) | 1.10 |  | STAT1 | MA0036.3<br>(GATA2) | 1.80 |
| — | Sin3Ak | MA0148.4<br>(FOXA1) | 0.98 |  | **TCF12** | MA0036.3<br>(GATA2) | **1.70** |
|  | ATF3 | MA0148.4<br>(FOXA1) | 0.96 |  | THAP1 | MA0036.3<br>(GATA2) | 1.50 |
|  | GABP | MA0148.4<br>(FOXA1) | 0.92 |  | Nrf1 | MA0036.3<br>(GATA2) | 1.30 |
| — | NRSF | MA0148.4<br>(FOXA1) | 0.85 |  | ZBTB7A | MA0036.3<br>(GATA2) | 1.10 |
|  | SP2 | MA0148.4<br>(FOXA1) | 0.57 |  | SRF | MA0036.3<br>(GATA2) | 0.95 |
|  | FOSL2 | MA0148.4<br>(FOXA1) | 0.57 |  | ATF3 | MA0036.3<br>(GATA2) | 0.90 |
| — | Rad21 | MA0148.4<br>(FOXA1) | 0.40 |  |  |  |  |

**Fig. 8:**
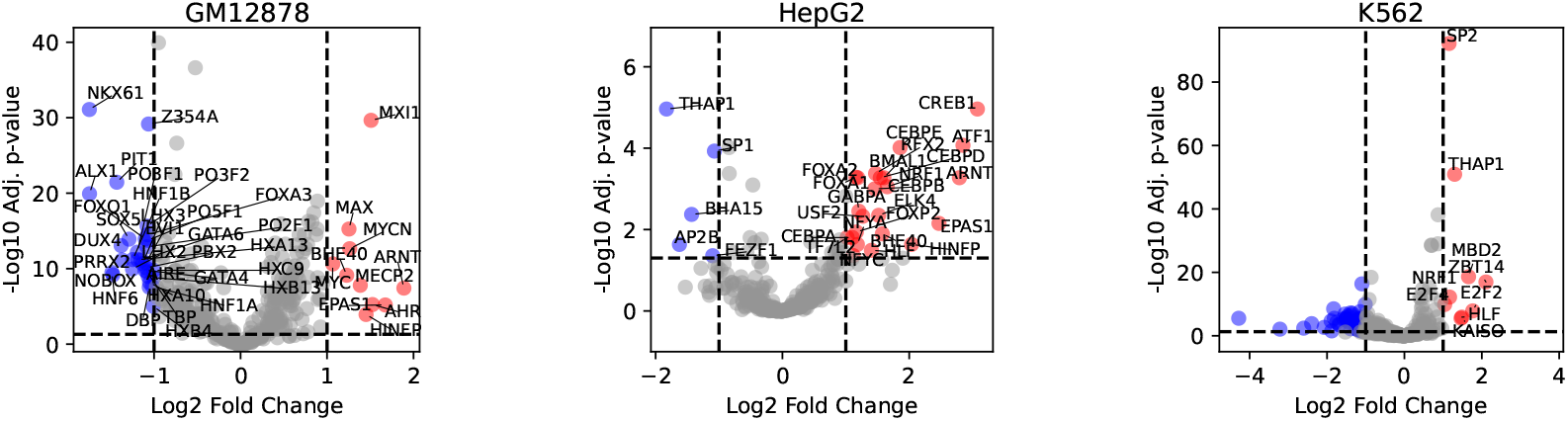
Motif enrichment analysis of demethylation-sensitive loci. Volcano plots showing enriched TF motifs identified by FIMO analysis on sites where demethylation increases TF binding (red) or depleted (blue) across three cell lines: GM12878, HepG2, and K562. The *x*-axis represents the log_2_ fold change in motif occurrence between methylation-sensitive loci and the remaining genomic background, while the *y*-axis shows the *−*log_10_ adjusted *p*-value. Dashed lines indicate significance thresholds.

**Fig. 9:**
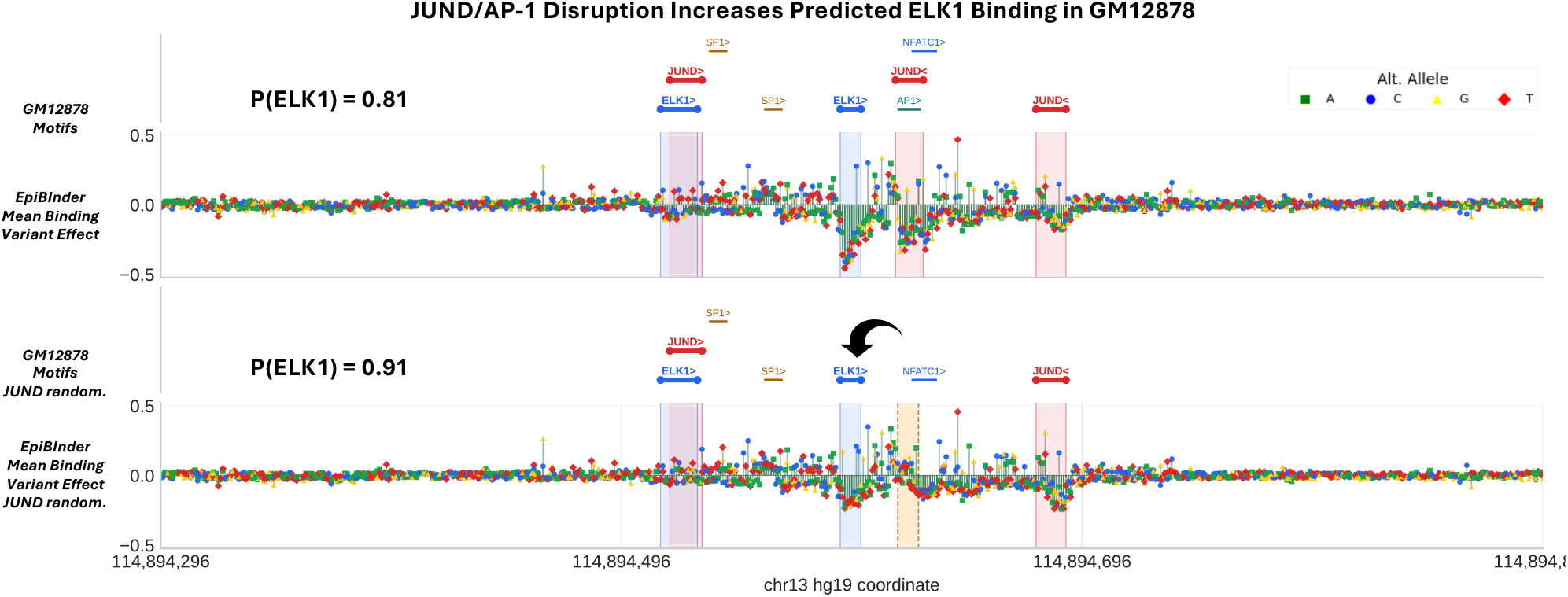
Randomizing a nearby JUND/AP-1 motif increases predicted ELK1 binding in GM12878. Single-nucleotide mutagenesis at a GM12878 regulatory sequence shows increased predicted ELK1 binding after disruption of a nearby JUND/AP-1 motif. The top track shows the native sequence context, with active-label motifs annotated by FIMO and JUND/ELK1 sites highlighted. The bottom track shows the same locus after randomizing the nearby JUND/AP-1 motif. The randomized interval is marked in the lower mutagenesis panel. Model-predicted ELK1 binding increases from 0.818 to 0.917 after JUND motif randomization, corresponding to Δ_*bind*_ = +0.1.

**Fig. 10:**
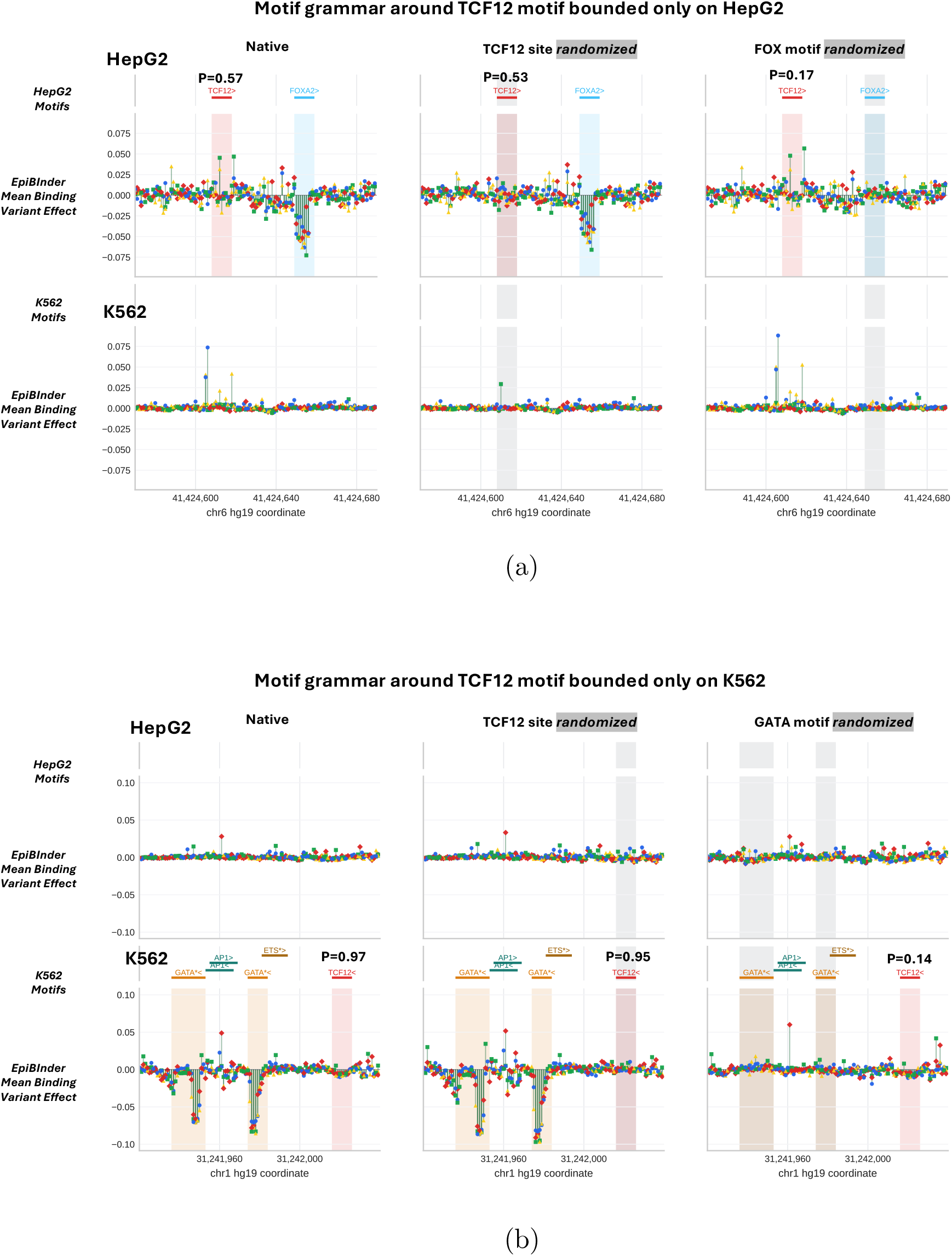
Example of cell-type-specific regulatory context underlying TCF12 binding. **(a)** A TCF12-binding locus at chr6:41424570– 41424690, where *EpiBinder* predicts TCF12 occupancy in HepG2, but not in K562, and the local mutagenesis profile assigns high sensitivity to a nearby FOX-associated motif. In silico randomization of this motif decreases the predicted TCF12 binding probability from 0.57 to 0.17 (Δ_bind_ = *−*0.40). **(b)** K562 locus at chr1:31241920–31242040, where alternatively *EpiBinder* predicts TCF12 occupancy in K562, but not in HepG2, with sensitivity this time concentrated on nearby GATA-associated motifs. Randomization of these motifs decreases the predicted TCF12 binding probability from 0.97 to 0.14 (Δ_bind_ = *−*0.83).

Finally, we extend the analysis of model-derived patterns revealing widespread contextual associations from GM12878 (Section 3.3) to HepG2 and K562, shown here in Fig. 11 and Fig. 12, respectively.

**Fig. 11:**
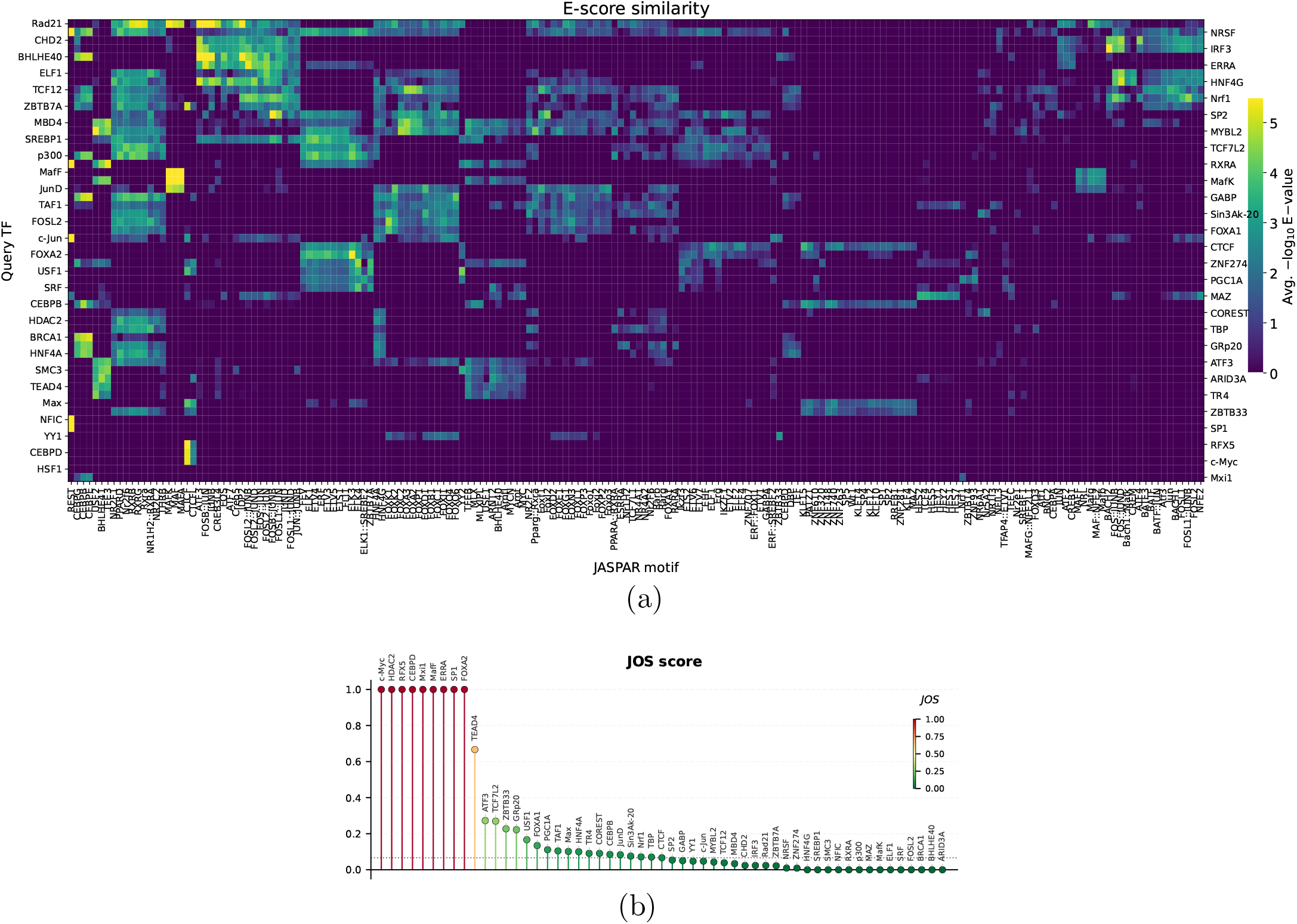
Model-derived patterns reveal widespread contextual associations (HepG2 cell-line). **(a)** Motif similarity between model-derived patterns and the JASPAR database reported as the Tomtom similarity score (*−* log_10_ of the E-value). Bright colors correspond to stronger similarity (lower E-values), while darker colors indicate low similarity. **(b)** Jaccard Overlap Score (JOS) summarizing diversity in the association for each TF. The score ranges from 0 (patterns exhibit wide associations with diverse motifs) to 1 (patterns converge on a single canonical motif).

**Fig. 12:**
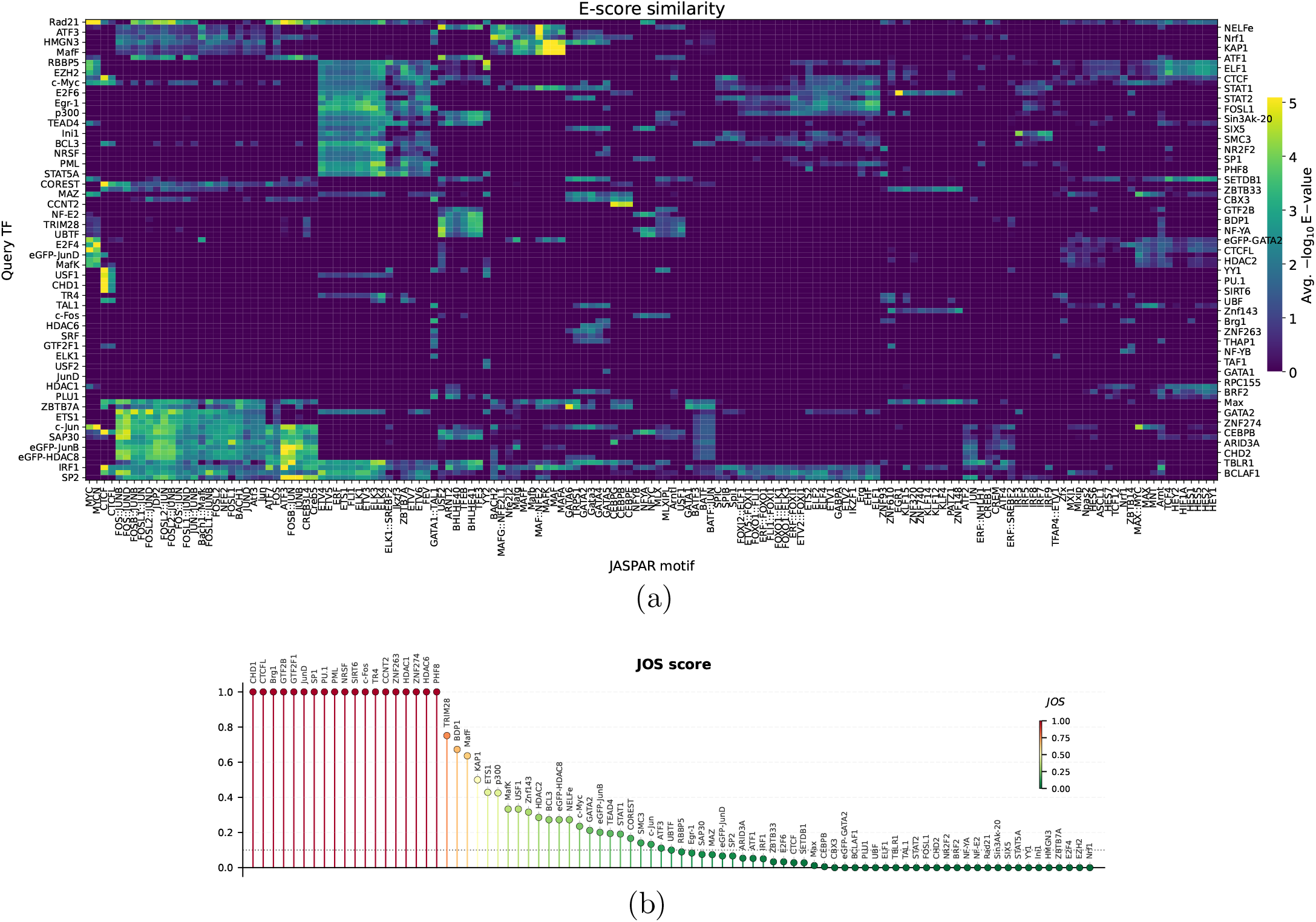
Model-derived patterns reveal widespread contextual associations (K562 cell-line). **(a)** Motif similarity between model-derived patterns and the JASPAR database reported as the Tomtom similarity score (*−* log_10_ of the E-value). Bright colors correspond to stronger similarity (lower E-values), while darker colors indicate low similarity. **(b)** Jaccard Overlap Score (JOS) summarizing diversity in the association for each TF. The score ranges from 0 (patterns exhibit wide associations with diverse motifs) to 1 (patterns converge on a single canonical motif).

## References

1. Vikram Agarwal, Fumitaka Inoue, Max Schubach, Dmitry Penzar, Beth K Martin, Pyaree Mohan Dash, Pia Keukeleire, Zicong Zhang, Ajuni Sohota, Jingjing Zhao, et al. Massively parallel characterization of transcriptional regulatory elements. Nature, 639(8054):411–420, 2025.

2. Babak Alipanahi, Andrew Delong, Matthew T Weirauch, and Brendan J Frey. Predicting the sequence specificities of dna-and rna-binding proteins by deep learning. Nature biotechnology, 33(8):831–838, 2015.

3. Christof Angermueller, Heather J Lee, Wolf Reik, and Oliver Stegle. Deepcpg: accurate prediction of single-cell dna methylation states using deep learning. Genome biology, 18:1–13, 2017.

4. Žiga Avsec, Vikram Agarwal, Daniel Visentin, Joseph R Ledsam, Agnieszka Grabska-Barwinska, Kyle R Taylor, Yannis Assael, John Jumper, Pushmeet Kohli, and David R Kelley. Effective gene expression prediction from sequence by integrating long-range interactions. Nature methods, 18(10):1196–1203, 2021.

5. Žiga Avsec, Natasha Latysheva, Jun Cheng, Guido Novati, Kyle R Taylor, Tom Ward, Clare Bycroft, Lauren Nicolaisen, Eirini Arvaniti, Joshua Pan, et al. Alphagenome: advancing regulatory variant effect prediction with a unified dna sequence model. bioRxiv, pages 2025–06, 2025.

6. Swneke D Bailey, Xiaoyang Zhang, Kinjal Desai, Malika Aid, Olivia Corradin, Richard Cowper-Sal·lari Batool Akhtar-Zaidi, Peter C Scacheri, Benjamin Haibe-Kains, and Mathieu Lupien. Znf143 provides sequence specificity to secure chromatin interactions at gene promoters. Nature communications, 6(1):6186, 2015.

7. Timothy L Bailey, Mikael Boden, Fabian A Buske, Martin Frith, Charles E Grant, Luca Clementi, Jingyuan Ren, Wilfred W Li, and William S Noble. Meme suite: tools for motif discovery and searching. Nucleic acids research, 37(suppl 2):W202–W208, 2009.

8. Bradley E Bernstein, John A Stamatoyannopoulos, Joseph F Costello, Bing Ren, Aleksandar Milosavljevic, Alexander Meissner, Manolis Kellis, Marco A Marra, Arthur L Beaudet, Joseph R Ecker, et al. The nih roadmap epigenomics mapping consortium. Nature biotechnology, 28(10):1045–1048, 2010.

9. Lu Chai, Jie Gao, Zihan Li, Hao Sun, Junjie Liu, Yong Wang, and Lirong Zhang. Predicting ctcf cell type active binding sites in human genome. Scientific Reports, 14(1):31744, 2024.

10. Jean-Thomas Clément and Qiao Li. Cistromic insight into the association of retinoid x receptor with myod and ctcf in proliferating myoblasts. Scientific Reports, 15(1):38919, 2025.

11. Hugo Dalla-Torre, Liam Gonzalez, Javier Mendoza-Revilla, Nicolas Lopez Carranza, Adam Henryk Grzywaczewski, Francesco Oteri, Christian Dallago, Evan Trop, Bernardo P de Almeida, Hassan Sirelkhatim, et al. The nucleotide transformer: Building and evaluating robust foundation models for human genomics. BioRxiv, pages 2023–01, 2023.

12. Iain F Davidson and Jan-Michael Peters. Genome folding through loop extrusion by smc complexes. Nature reviews Molecular cell biology, 22(7):445–464, 2021.

13. Lucas Paulo de Lima Camillo, Raghav Sehgal, Jenel Armstrong, Albert Tzongyang Higgins-Chen, Steve Horvath, and Bo Wang. Cpgpt: a foundation model for dna methylation. bioRxiv, pages 2024–10, 2024.

14. Natalie de Souza. The encode project. Nature methods, 9(11):1046–1046, 2012.

15. Christopher L Frank, Dinesh Manandhar, Raluca Gordân, and Gregory E Crawford. Hdac inhibitors cause site-specific chromatin remodeling at pu. 1-bound enhancers in k562 cells. Epigenetics & chromatin, 9(1):15, 2016.

16. Geoff Fudenberg, David R Kelley, and Katherine S Pollard. Predicting 3d genome folding from dna sequence with akita. Nature methods, 17(11):1111–1117, 2020.

17. Tianshun Gao and Jiang Qian. Enhanceratlas 2.0: an updated resource with enhancer annotation in 586 tissue/cell types across nine species. Nucleic acids research, 48(D1):D58–D64, 2020.

18. GENA LM. GENA LM. https://github.com/AIRI-Institute/GENA_LM/tree/main/downstream_tasks/DeepSea, 2024.

19. Charles E Grant, Timothy L Bailey, and William Stafford Noble. Fimo: scanning for occurrences of a given motif. Bioinformatics, 27(7):1017–1018, 2011.

20. Sven Heinz, Christopher Benner, Nathanael Spann, Eric Bertolino, Yin C Lin, Peter Laslo, Jason X Cheng, Cornelis Murre, Harinder Singh, and Christopher K Glass. Simple combinations of lineage-determining transcription factors prime cis-regulatory elements required for macrophage and b cell identities. Molecular cell, 38(4):576–589, 2010.

21. Aldo Hernandez-Corchado and Hamed S Najafabadi. Toward a base-resolution panorama of the in vivo impact of cytosine methylation on transcription factor binding. Genome Biology, 23(1):151, 2022.

22. Makiko Iwafuchi-Doi, Greg Donahue, Akshay Kakumanu, Jason A Watts, Shaun Mahony, B Franklin Pugh, Dolim Lee, Klaus H Kaestner, and Kenneth S Zaret. The pioneer transcription factor foxa maintains an accessible nucleosome configuration at enhancers for tissue-specific gene activation. Molecular cell, 62(1):79–91, 2016.

23. Yanrong Ji, Zhihan Zhou, Han Liu, and Ramana V Davuluri. Dnabert: pre-trained bidirectional encoder representations from transformers model for dna-language in genome. Bioinformatics, 37(15):2112–2120, 2021.

24. Alireza Karbalayghareh, Merve Sahin, and Christina S Leslie. Chromatin interaction–aware gene regulatory modeling with graph attention networks. Genome Research, 32(5):930–944, 2022.

25. Martin Kircher, Chenling Xiong, Beth Martin, Max Schubach, Fumitaka Inoue, Robert JA Bell, Joseph F Costello, Jay Shendure, and Nadav Ahituv. Saturation mutagenesis of twenty disease-associated regulatory elements at single base-pair resolution. Nature communications, 10(1):3583, 2019.

26. Sandy L Klemm, Zohar Shipony, and William J Greenleaf. Chromatin accessibility and the regulatory epigenome. Nature Reviews Genetics, 20(4):207–220, 2019.

27. Judith F Kribelbauer, Xiang-Jun Lu, Remo Rohs, Richard S Mann, and Harmen J Bussemaker. Toward a mechanistic understanding of dna methylation readout by transcription factors. Journal of molecular biology, 432(6):1801–1815, 2020.

28. Mengwei Li, Dong Zou, Zhaohua Li, Ran Gao, Jian Sang, Yuansheng Zhang, Rujiao Li, Lin Xia, Tao Zhang, Guangyi Niu, Yiming Bao, and Zhang Zhang. EWAS atlas: a curated knowledgebase of epigenome-wide association studies. Nucleic Acids Res., 47(Database-Issue):D983–D988, 2019.

29. Yan Li, Judith HI Haarhuis, Ángela Sedeño Cacciatore, Roel Oldenkamp, Marjon S van Ruiten, Laureen Willems, Hans Teunissen, Kyle W Muir, Elzo de Wit, Benjamin D Rowland, et al. The structural basis for cohesin–ctcf-anchored loops. Nature, 578(7795):472–476, 2020.

30. Jakub Lipinski. Build DeepSEA training dataset. https://github.com/jakublipinski/build-deepsea-training-dataset, 2024.

31. Yuting Liu, Xin Wan, Hu Li, Yingxi Chen, Xiaodi Hu, Hebing Chen, Dahai Zhu, Cheng Li, and Yong Zhang. Ctcf coordinates cell fate specification via orchestrating regulatory hubs with pioneer transcription factors. Cell reports, 42(10), 2023.

32. Scott M Lundberg and Su-In Lee. A unified approach to Llion Jones, Aidan N Gomez, L-ukasz Kaiser, and Illia interpreting model predictions. Advances in neural information processing systems, 30, 2017.

33. Bethany J Madison, Kathleen A Clark, Niraja Bhachech, Peter C Hollenhorst, Barbara J Graves, and Simon L Currie. Electrostatic repulsion causes anticooperative dna binding between tumor suppressor ets transcription factors and jun– fos at composite dna sites. Journal of Biological Chemistry, 293(48):18624–18635, 2018.

34. Eric Nguyen, Michael Poli, Marjan Faizi, Armin Thomas, Michael Wornow, Callum Birch-Sykes, Stefano Massaroli, Aman Patel, Clayton Rabideau, Yoshua Bengio, et al. Hyenadna: Long-range genomic sequence modeling at single nucleotide resolution. Advances in neural information processing systems, 36, 2024.

35. Sierra S Nishizaki and Alan P Boyle. Semplme: a tool for integrating dna methylation effects in transcription factor binding affinity predictions. BMC bioinformatics, 23(1):317, 2022.

36. Zaneta Odrowaz and Andrew D Sharrocks. Elk1 uses different dna binding modes to regulate functionally distinct classes of target genes. PLoS genetics, 8(5):e1002694, 2012.

37. E Christopher Partridge, Surya B Chhetri, Jeremy W Prokop, Ryne C Ramaker, Camden S Jansen, Say-Tar Goh, Mark Mackiewicz, Kimberly M Newberry, Laurel A Brandsmeier, Sarah K Meadows, et al. Occupancy maps of 208 chromatin-associated proteins in one human cell type. Nature, 583(7818):720–728, 2020.

38. Sachin Pundhir, Felicia Kathrine Bratt Lauridsen, Mikkel Bruhn Schuster, Janus Schou Jakobsen, Ying Ge, Erwin Marten Schoof, Nicolas Rapin, Johannes Waage, Marie Sigurd Hasemann, and Bo Torben Porse. Enhancer and transcription factor dynamics during myeloid differentiation reveal an early differentiation block in cebpa null progenitors. Cell reports, 23(9):2744–2757, 2018.

39. Yitzhak Reizel, Ashleigh Morgan, Long Gao, Jonathan Schug, Sarmistha Mukherjee, Meilín FernÁndez García, Greg Donahue, Joseph A Baur, Kenneth S Zaret, and Klaus H Kaestner. Foxa-dependent demethylation of dna initiates epigenetic memory of cellular identity. Developmental Cell, 56(5):602–612, 2021.

40. Albin Sandelin, Wynand Alkema, Pär Engström, Wyeth W Wasserman, and Boris Lenhard. Jaspar: an open-access database for eukaryotic transcription factor binding profiles. Nucleic acids research, 32(suppl 1):D91–D94, 2004.

41. Nagarathinam Selvaraj, Justin A Budka, Mary W Ferris, Joshua P Plotnik, and Peter C Hollenhorst. Extracellular signal-regulated kinase signaling regulates the opposing roles of jun family transcription factors at ets/ap-1 sites and in cell migration. Molecular and Cellular Biology, 35(1):88–100, 2015.

42. Dustin Shigaki, Orit Adato, Aashish N Adhikari, Shengcheng Dong, Alex Hawkins-Hooker, Fumitaka Inoue, Tamar Juven-Gershon, Henry Kenlay, Beth Martin, Ayoti Patra, et al. Integration of multiple epigenomic marks improves prediction of variant impact in saturation mutagenesis reporter assay. Human mutation, 40(9):1280–1291, 2019.

43. Avanti Shrikumar, Katherine Tian, Žiga Avsec, Anna Shcherbina, Abhimanyu Banerjee, Mahfuza Sharmin, Surag Nair, and Anshul Kundaje. Technical note on transcription factor motif discovery from importance scores (tf-modisco) version 0.5. 6.5. arXiv preprint arXiv:1811.00416, 2018.

44. Ashish Vaswani, Noam Shazeer, Niki Parmar, Jakob Uszkoreit Polosukhin. Attention is all you need. Advances in neural information processing systems, 30, 2017.

45. Timothy Warwick, Marcel H Schulz, Ralf Gilsbach, Ralf P Brandes, and Sabine Seuter. Nuclear receptor activation shapes spatial genome organization essential for gene expression control: Lessons learned from the vitamin d receptor. Nucleic acids research, 50(7):3745–3763, 2022.

46. Yumi Minyi Yao, Irina Miodownik, Michael P O’Hagan, Muhammad Jbara, and Ariel Afek. Deciphering the dynamic code: Dna recognition by transcription factors in the ever-changing genome. Transcription, 15(3-5):114–138, 2024.

47. Kejun Ying, Jinyeop Song, Haotian Cui, Yikun Zhang, Siyuan Li, Xingyu Chen, Hanna Liu, Alec Eames, Daniel L McCartney, Riccardo E Marioni, et al. Methylgpt: a foundation model for the dna methylome. bioRxiv, pages 2024–10, 2024.

48. Manzil Zaheer, Guru Guruganesh, Kumar Avinava Dubey, Joshua Ainslie, Chris Alberti, Santiago Ontanon, Philip Pham, Anirudh Ravula, Qifan Wang, Li Yang, et al. Big bird: Transformers for longer sequences. Advances in neural information processing systems, 33:17283–17297, 2020.

49. Jian Zhou and Olga G Troyanskaya. Predicting effects of noncoding variants with deep learning–based sequence model. Nature methods, 12(10):931–934, 2015.

50. Q Zhou, M Yu, R Tirado-Magallanes, B Li, L Kong, M Guo, ZH Tan, S Lee, L Chai, A Numata, et al. Znf143 mediates ctcf-bound promoter-enhancer loops required for murine hematopoietic stem and progenitor cell function. nat. commun. 12, 43, 2021.

51. Zhihan Zhou, Yanrong Ji, Weijian Li, Pratik Dutta, Ramana Davuluri, and Han Liu. Dnabert-2: Efficient foundation model and benchmark for multi-species genome. arXiv preprint arXiv:2306.15006, 2023.

